# Dynamic states of cervical epithelia during pregnancy and epithelial barrier disruption

**DOI:** 10.1101/2022.07.26.501609

**Authors:** Anne Cooley, ShanmugaPriyaa Madhukaran, Elizabeth Stroebele, Mariano Colon Caraballo, Lei Wang, Gary C. Hon, Mala Mahendroo

## Abstract

The cervical epithelium undergoes continuous changes in proliferation, differentiation, and function that are critical before pregnancy to ensure fertility and during pregnancy to provide a physical and immunoprotective barrier for pregnancy maintenance. Barrier disruption can lead to the ascension of pathogens that elicit inflammatory responses and preterm birth. Here, we identify cervical epithelial subtypes in nonpregnant, pregnant, and in-labor mice using single-cell transcriptome and spatial analysis. We identify heterogeneous subpopulations of epithelia displaying spatial and temporal specificity. Notably, two goblet cell subtypes with distinct transcriptional programs and mucosal networks were dominant in pregnancy. Untimely basal cell proliferation and goblet cells with diminished mucosal integrity characterize barrier dysfunction in mice lacking hyaluronan. These data demonstrate how the cervical epithelium undergoes continuous remodeling to maintain dynamic states of homeostasis in pregnancy and labor, and provide a framework to understand perturbations in epithelial health and host-microbe interactions that increase the risk of premature birth.

## INTRODUCTION

Cervical epithelial cells have diverse roles during the nonpregnant (NP) reproductive cycle, throughout pregnancy, and parturition. These cells ensure fertility, provide a physical and immunological barrier to prevent the ascension of pathogens to the upper reproductive tract, and elicit signals that kill non-commensal pathogens(Wira et al., 2015). Cervical epithelia carry out these functions in the context of a dynamic tissue that undergoes continuous remodeling before, during, and after pregnancy (Nallasamy and Mahendroo, 2017).

Driven by the ovarian steroid hormones progesterone (P4) and estrogen (E2), cervical epithelia undergo numerous structural, morphological and functional changes throughout the NP reproductive cycle. These hormones regulate proliferation, differentiation, mucus secretion, and the ability of epithelial cells to respond to pathogenic microbes (Wira et al., 2015). As a key component of the mucosal immune system, vaginal and cervical epithelia produce mucins, cytokines, antimicrobial molecules, and transport immunoglobulins essential to protect the female reproductive tract from the invasion of pathogens (Sugiyama et al., 2021; Wira et al., 2005). The mucosal environment is finely balanced to provide pathogen surveillance and facilitate the migration of sperm from the cervicovaginal canal to the fallopian tube for fertilization (Barrios De Tomasi et al., 2019; Givan et al., 1997; Goode et al., 2014). Disruptions in the epithelial barrier can lead to sexually transmitted infections and/or infertility (McShane et al., 2021) (Anahtar et al., 2018).

Numerous studies report changes in the levels of cytokines, antimicrobial molecules, and immunoglobulins in cervicovaginal fluids of pregnant women relative to NP women, suggesting an alteration in epithelial subtypes or responses in the unique hormonal milieu of pregnancy (Anderson et al., 2013; Hughes et al., 2016; Kutteh and Franklin, 2001; Walter et al., 2011). During pregnancy, disruptions in the epithelial barrier or increased mucus permeability are associated with increased susceptibility to ascending infections and preterm birth in women. In mice, disruptions of the epithelial barrier due to loss of hyaluronan, chemical disruption of loss of the gel-forming mucin, Muc5b increase rates of ascending infection mediated preterm birth (Akgul et al., 2014; Burris et al., 2020; Critchfield et al., 2013; Hezelgrave et al., 2020; Kyrgiou et al., 2006; Lacroix et al., 2022; Pavlidis et al., 2020; Smith-Dupont et al., 2017). Despite many efforts, effective therapies for prevention are lacking due to an incomplete understanding of the specialized epithelial subtypes in the cervix, their functions and their regulation throughout pregnancy and parturition. Hence, understanding the dynamic cell state changes during pregnancy could yield insights into mechanisms of normal and preterm cervical remodeling.The cervical epithelia is divided into endocervical and ectocervical regions (**Fig 1A**). In mice, both regions contain stratified squamous epithelium and lack glands (Cunha et al., 2019; Fishbeck and Sebastiani, 2015; Leppi and John Leppi, 1964; Mehta et al., 2016; 2011). The mouse endocervix also contains columnar epithelium that vary during development and the estrus cycle (Kurita et al., 2001). The human endocervix consists of columnar and squamous epithelium and contains glands. The ectocervix is solely stratified squamous epithelium (Cunha et al., 2017; Singer, 1977). Despite potential differences in endocervical epithelial subtypes between mouse and human, both humans and rodents produce a similar mucus network and are fortified with a similar arsenal of protective factors (Gipson et al., 1997; Portal et al., 2017; Richardson et al., 1993).

**Figure 1:**
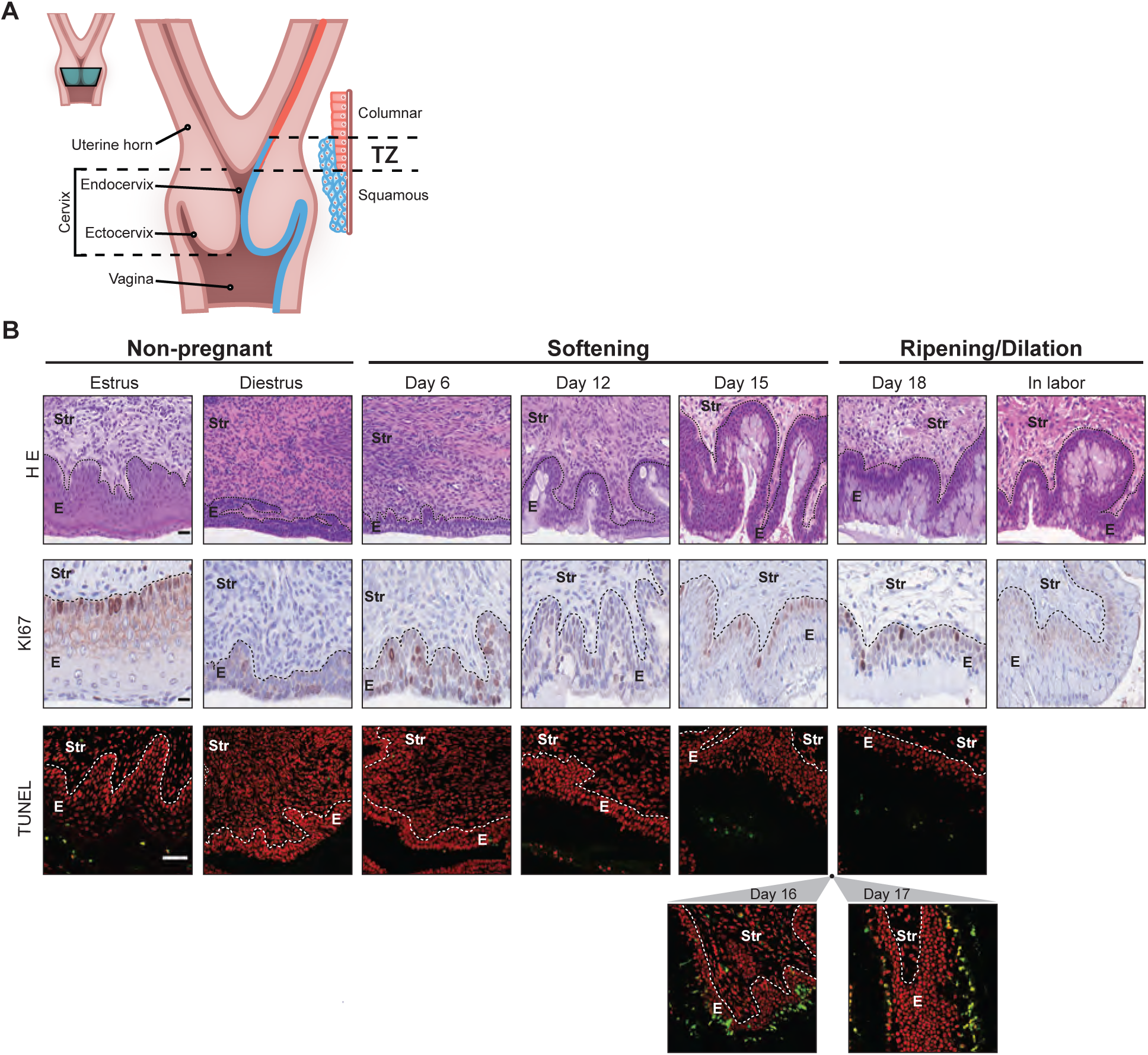
Morphological changes in cervical epithelia during pregnancy. A. Schematic longitudinal section of mouse uterus and cervix. Blue: stratified squamous epithelia of endo- and ectocervix. Red: columnar epithelium of uterus. Transition zone (TZ): Boundary comprising both types of epithelia. Green boxed area: Indicates cervical region below TZ collected for single cell libraries. B. Morphological changes in cervical epithelia in non-pregnant, pregnant (D6, D12, D15 and D18), and in labor mice. Shown are H&E staining (top), Ki67 immunostaining for proliferation (middle), and TUNEL staining for cell death (bottom) (red: nuclei, green: TUNEL^+^). E: Epithelia; Str: Stroma. Scale bar 50 µm, Representative images from three independent samples per each group.

The focus of this study is to identify cervical epithelial subtypes in the cervix of nonpregnant and pregnant mice and to evaluate their spatial location in the endo- and ectocervix. Here, we define transcriptional cell states of the cervical epithelia at four gestational time points over the 19-day mouse pregnancy relative to the cervix of in labor (IL) and NP mice. Using single cell genomics and spatial analysis, we define dynamic changes in cell states that begin in early pregnancy. We identify a misregulation in cell state transition associated with a mouse model of epithelial barrier disruption with increased susceptibility to ascending infection-induced preterm birth.

## RESULTS

### Morphological changes in cervical epithelia during pregnancy

We performed time course imaging analysis to examine the dynamic morphological changes during pregnancy. The multi-layered stratified epithelium undergoes marked changes in proliferation, differentiation, and apoptosis throughout the NP estrus cycle and 19-day pregnancy (**Fig 1B** **and Fig S1**). Most notable during pregnancy is the increase in secretory epithelia that peaks in cell size and cell number on gestation day 18 (GD18) and IL, as seen by H&E staining (**Fig 1B**). Proliferative (Ki67^+^) cells are present in all layers of epithelia on GD6 and 12, yet are confined primarily to the basal layers on GD15, GD18, and IL. Cell death is markedly induced in the luminal layer of cells in the endocervix on days 16 and 17 (TUNEL^+^ cells in green). During pregnancy, cell morphology and proliferation are similar between endo- and ectocervix while cell death occurs primarily in the endocervix. In contrast, during the NP diestrus stage, secretory epithelia are abundant in the ecto- but not endocervix (**Fig S1)**. These observations highlight the dynamic structural and functional reorganization of cervical epithelia during pregnancy.

### Dynamic transcriptional and epigenetic states of cervical epithelia during pregnancy

To spatially define the endocervix, the uterine epithelia and boundary between endocervix and uterus (Cunha et al., 2017), we performed immunofluorescence imaging for markers of squamous basal cells (keratin 5) and columnar epithelia (keratin 8) in the nonpregnant proliferative (estrus) and secretory (diestrus) stages (**Supplementary Figures 2 and 3**). The endocervix region was defined by the absence of glands and the presence of squamous Krt5+ cells (estrus **Fig S2)** or Krt5+ and Krt8+ cells (diestrus, **Fig S3**). The uterine region was defined by the presence of Krt8+ cells in both luminal and glandular epithelium. The transition zone (TZ) where columnar and squamous epithelia overlap was identified just below the uterine region, thus defining the boundary between uterus and endocervix as previously described (Fishbeck and Sebastiani, 2015; Kurita et al., 2001).

**Figure 2:**
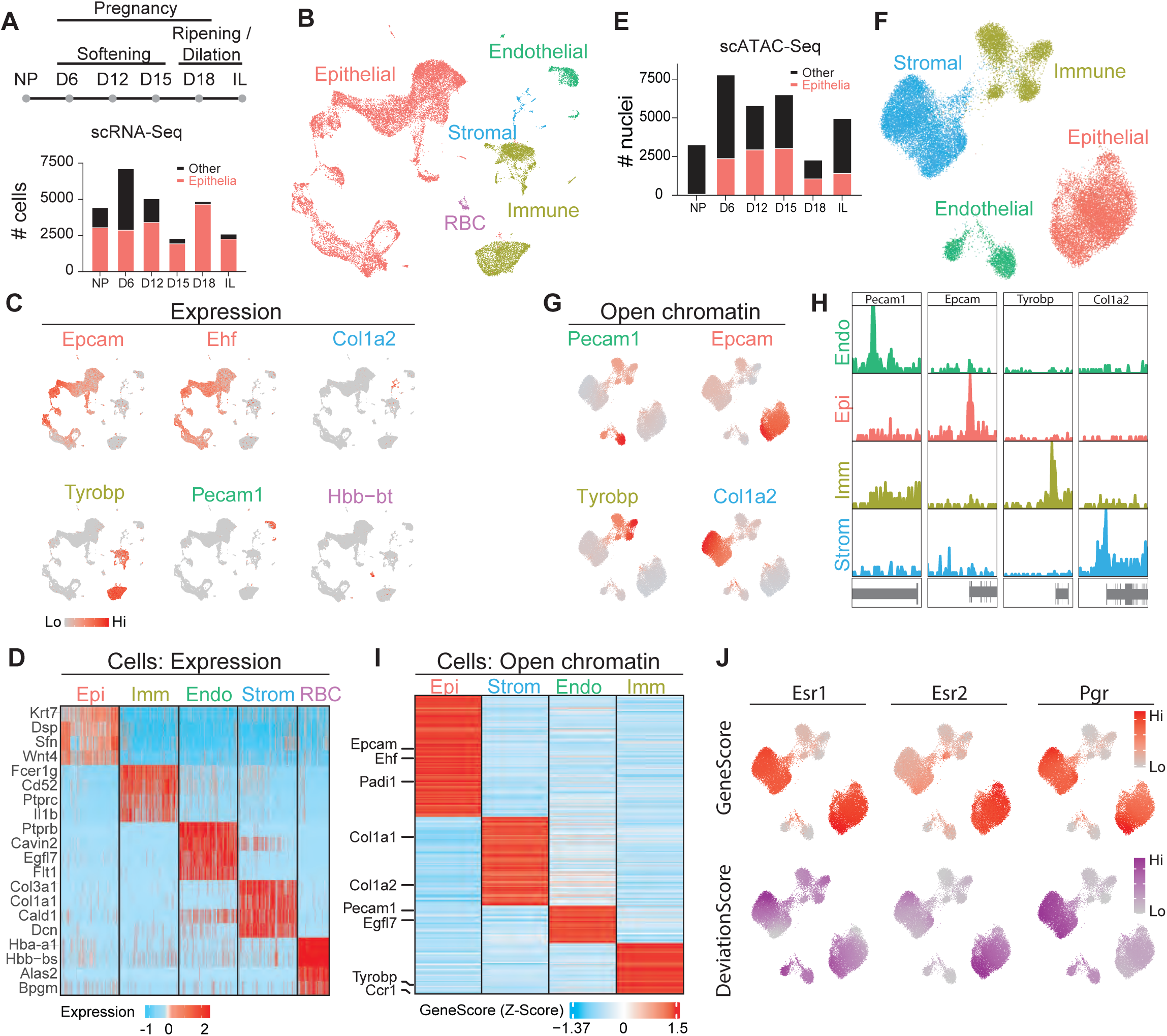
Dynamic transcriptional and epigenetic states of cervical epithelia during pregnancy. A. (top) Design of scRNA-Seq experiments (gray circles: sequencing time points). (bottom) Distribution of epithelia and non-epithelial cells captured per time point. B. UMAP visualization of all cells captured from all time points of scRNA-Seq. C. Feature plots indicate the expression of genes used to identify different cell types. Epcam and Ehf: epithelia; Col1a2: stroma; Tyrobp: immune; Pecam1: endothelia, and Hbb-bt: red blood cells. D. Heatmap showing the expression of cell-type specific genes. E. Distribution of epithelia and non-epithelial cells captured per time point of scATAC-Seq. F. UMAP visualization of all cells captured from all time points of scATAC-Seq. G. Feature plots indicate the open chromatin status of genes used to identify different cell types. H. Genome browser snapshots of open chromatin status for the genes in (G), for each cell type. I. Heatmap showing the cell-type specific open chromatin status of genes. J. (top) Feature plot of open chromatin status for Esr1, Esr2, and Pgr. (bottom) Motif deviation scores for these transcription factors.

To define the dynamic transcriptional states of cervical epithelia, we performed time-course single-cell RNA sequencing. Mouse cervical tissue was separated from the uterus just below the TZ (**Fig 1A**, green box) and harvested for scRNA-Seq library preparation using cell digestion methods optimized to ensure epithelial cell viability. The NP library consists of cells in the secretory (diestrus) and proliferative (estrus) stages of the cycle. Time points in pregnancy were selected to span the phases of cervical softening, ripening, and day 19 in labor after delivery of 1-2 pups (**Fig 2A**). To ensure sufficient cell numbers, cervices were pooled from the indicated number of mice: NP (n=9; 7 diestrus and 2 estrus), GD6 (n= 7), GD12 (n= 2), GD15 (n=3), GD18 (n=2), and IL (n=2). After stringent quality control filters (see Methods), we recovered 26,493 high-quality cells across the six time points. Epithelial cells were the main cell type recovered at each time point while the number of stromal cells captured was low relative to their abundance in tissue (**Fig 2A**), as observed in another study (Garcia-Flores et al., 2022). Clustering of all time points showed five distinct cell types that were captured from the cervical tissue: epithelial (Epcam^+^, Ehf^+^), immune (Tyrobp^+^), endothelial (Pecam1^+^), stromal (Col1a2^+^), and red blood cells (Hbb-bt^+^) (**Fig 2B and C**) (Becht et al., 2018). Additional cell type-specific genes also confirm the identity of the cell types (**Fig 2D**). Since epithelial cells were the dominant cell type captured at each time point, we focused downstream analyses on characterizing how epithelial cell states change throughout pregnancy.

To define the epigenetic states of cervical cell types, we also performed single-cell ATAC-Seq (Satpathy et al., 2019) throughout pregnancy time points. We recovered 30,758 nuclei (**Fig 2E**) that represent multiple cell types of the cervix with distinct open chromatin profiles (**Fig 2F**). We annotated each cluster by examining the open chromatin status for cell-type specifically expressed genes: epithelial (Epcam^+^), immune (Tyrobp^+^), endothelial (Pecam1^+^), and stromal (Col1a2^+^) (**Fig 2G-H**). The open chromatin status of additional cell-type specifically expressed genes also confirm these annotations (**Fig 2I**). Finally, by examining the enrichment of motifs at open chromatin peaks, we can identify transcription factors with potential roles in regulating specific cell types (Granja et al., 2021) during cervical remodeling. This analysis suggests divergent roles for well-known nuclear hormone receptors. For example, estrogen receptor 1 (Esr1) and progesterone receptor (Pgr) are active (gene expression and enriched open chromatin for the binding site) in both stromal and epithelial cells, while Esr2 is more active in epithelial cells (**Fig 2J**).

### Shifts in epithelial subtypes and proliferation in early softening

Previous bulk transcriptomic and proteomic studies suggest a marked change in epithelial cell phenotype and function during early cervical softening relative to NP (Nallasamy et al., 2021). To define the epithelial subtypes and cell state transitions induced in the early softening period, we compared epithelial subtypes in NP and GD6. The squamous epithelia are subdivided into basal, luminal (defined as a non-secretory luminal cell), and secretory luminal cells (defined as cells expressing mucin genes).

We identified basal, luminal, and secretory clusters in NP and GD6 samples (**Fig 3B and D**) with a similar distribution of cell types (**Fig 3F**). Notably, we observed an increase in proliferating (Ki67^+^) non-basal cells on GD6 compared to NP (**Fig 3C, 3E, 3G**). We also observed changes in luminal markers over time, with Dsg1a specifically marking non-secretory luminal cells in NP and Krt12 specifically marking non-secretory luminal cells in D6 (**Fig 3C, 3E**). Krt12 has not previously been described in the cervix and is considered a marker of the differentiated epithelium of the cornea (Kasetti et al., 2016).

**Figure 3:**
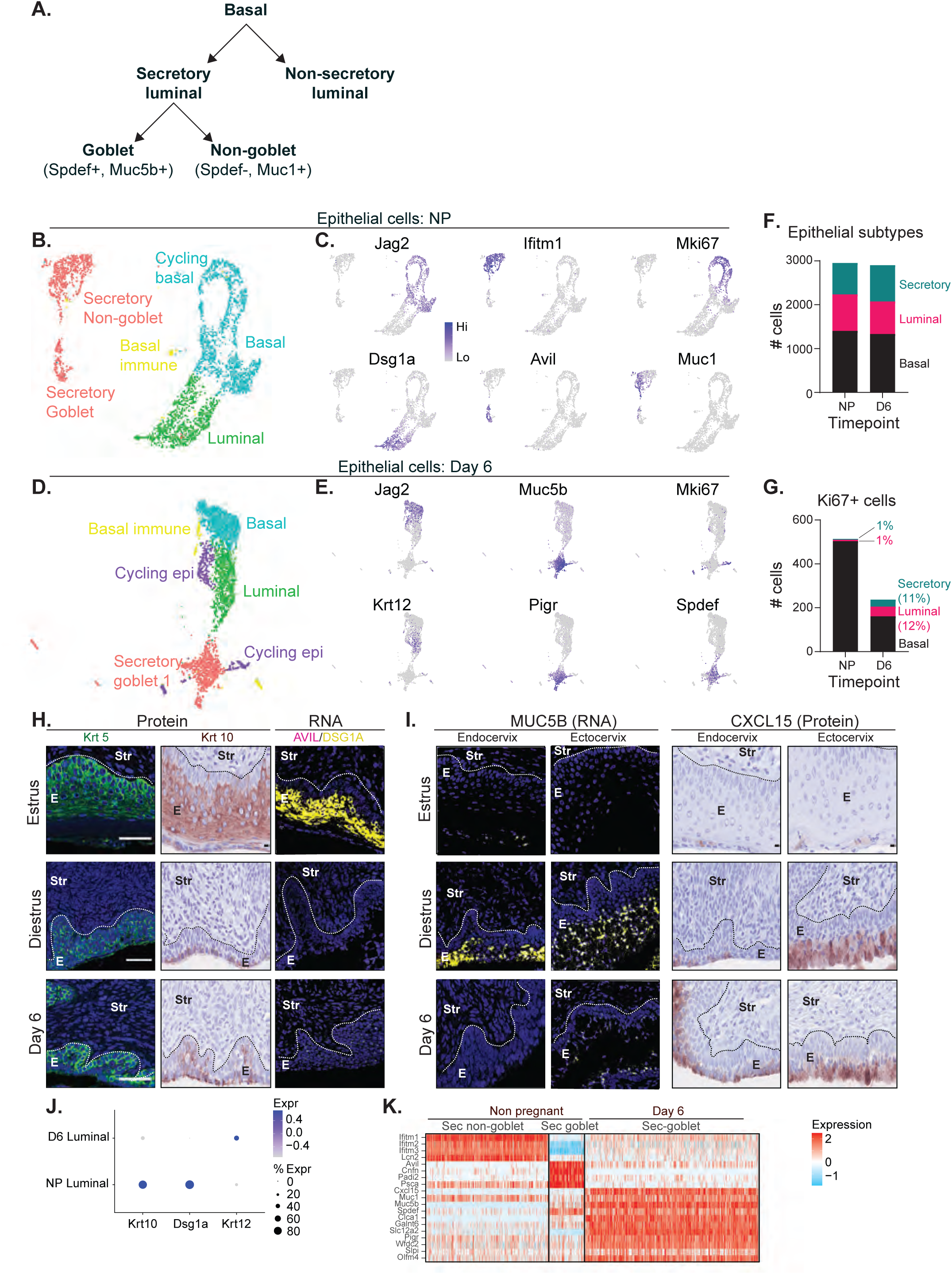
Shifts in epithelial subtypes and proliferation in early softening. A. Schematic of lineage relationships of cervical epithelial cell types. Basal cells give rise to luminal non-secretory and luminal secretory cells. Luminal secretory cells can mature into non-goblet (Spdef-Muc15^+^) or goblet cells (Spdef^+^, Muc5b^+^). B. UMAP visualization of epithelial cells from the non-pregnant time point with three major types of epithelia: basal, luminal, and secretory. C. Feature plots showing the expression of genes differentially expressed between the NP epithelia. Jag2: basal cells; Mki67: cycling basal cells; Dsg1a: luminal cells; Ifitm1, Avil, and Muc1: secretory cells. D. UMAP visualization of epithelial cells from GD6. E. Feature plots showing the expression of genes differentially expressed between the different types of GD6 epithelia. Jag2: basal cells; Mki67: cycling basal cells; Krt12: luminal cells; Muc5b, Pigr, and Spdef: secretory cells. F. Bar chart quantifying epithelial subtypes in NP and GD6. G. Bar chart quantifying proliferating (Mki67^+^)epithelial cells in NP and D6. H. IF and IHC showing protein expression of Krt5 (green), and Krt10 (brown) in mouse endocervical epithelia from NP (estrus and diestrus) and GD6. Krt5: basal cells; Krt10: luminal epithelia. RNAscope analysis of Dsg1a (yellow) with Avil (pink) mRNA in mouse endocervical epithelia from NP and GD6. DAPI (blue)- nuclei. E: Epithelia; Str: Stroma.Scale bar 50 µm, objective lens 40x. Representative images from two independent samples per group. I. Expression of Muc5b (yellow) mRNA by RNAscope and Cxcl15 (brown) protein by IHC in mouse endo and ectocervical epithelia from NP (estrus and diestrus) and GD6. DAPI: nuclei. E: Epithelia; Str: Stroma. Representative images from three independent samples per group. J. Dot plot showing the expression of luminal markers. K. Heatmap highlighting the different genes expressed in secretory clusters from NP and D6.

In NP, we identified two secretory clusters expressing mucin 1 (Muc1) that were distinguished by the markers Ifitm1 and Avil (**Fig 3C**). In contrast, on GD6, we identified only one secretory population (Muc5b+, Pigr+). The GD6 secretory cluster and the NP Avil+ cluster expressed the transcription factor Spdef which is required for the differentiation of goblet cells, a specialized secretory cell that produces gel-forming mucins (Chen et al., 2009; Knoop and Newberry, 2018). Goblet cells have been described in the NP cervix (Portal et al., 2017). Thus we further subdivided the secretory clusters into Spdef^+^/Muc5b^+^ goblet cells or Spdef^-^/Muc1^+^ non-goblet secretory cells (**Fig3A).** Spatial analysis of protein or RNA expression for basal (Krt5^+^), luminal (Krt10^+^/Dsg1a^+^) and secretory (Avil^+^/Muc5b^+^/Cxcl15^+^) markers confirm a shift in luminal subtypes from NP to GD6 (endocervix, **Fig 3H, 3I)** and demonstrate similar patterns of expansion between the endo and ectocervix (**Supl Fig 5,6,7**). In the NP estrus stage, Krt10^+^/Dsg1a^+^ luminal cells are abundant (**Fig 3H**) while in the NP diestrus stage Avil^+^/Muc5b^+^/Cxcl15^+^ secretory luminal cells are abundant (**Fig 3I**). Avil^+^ secretory cells are visible in the endocervical luminal layers in diestrus, with fewer Avil^+^ cells in estrus and on GD6. Further, during NP- diestrus and GD6 we observe, goblet secretory cell expansion is similar in both the ectocervix and the endocervix as indicated by Muc5b and Cxcl15 expression (**Fig 3I** **and Sup** **Fig 7**). Cxcl15 is a chemokine expressed in numerous mucosal and endocrine organs and is a specific marker of glandular but not luminal epithelia of the uterus (Schmitz et al., 2007; Wang et al., 2018). The presence of distinct luminal subtypes in NP and GD6 is indicated in the dot plot (**Fig 3J**) and the secretory clusters in **Fig 3K**. Collectively, these data indicate two secretory populations in the NP cervix, one of which is a Avil^+^/Spdef^+^/Muc5b^+^/Muc1+ goblet cell and the other is (Ifitm1+, Muc1+) (**Fig 3K**). On GD6, a single secretory goblet cell population is identified that is transcriptionally distinct from the NP goblet cell.

### Distinct populations of secretory cells during pregnancy

To examine cellular dynamics during pregnancy, we focused on the 4 pregnancy time points between GD6 and GD18 (**Fig 4A**), particularly the secretory populations. Cluster analysis identifies two populations of Muc5b^+^/Spdef^+^ goblet cells: Goblet 1 cells express Pigr while Goblet 2 cells express Rbp2 (**Fig 4B**). Goblet 1 cells are most abundant in the scRNA-Seq library on GD6 and gradually decline towards GD18, while goblet 2 cells are rare in the earlier time points and peak in the GD18 library (**Fig 4C**). RNA velocity analysis suggests luminal cells give rise to goblet 2 cells but not goblet 1 cells (**Fig 4D**). Luminal and goblet 2 cells express Rbp2 (**Fig 4B**), while goblet 1 does not. RNA velocity analysis does not indicate that goblet 1 cells arise from any other sequenced cells. We also identified a minor secretory cluster with a secretory-like keratin expression pattern (Krt8/18) but lacking the goblet markers Spdef and Muc5b. This non-goblet secretory cluster also uniquely expresses Prap1, Napsa, C3, and Gpx3 (**Fig S8**).

**Figure 4:**
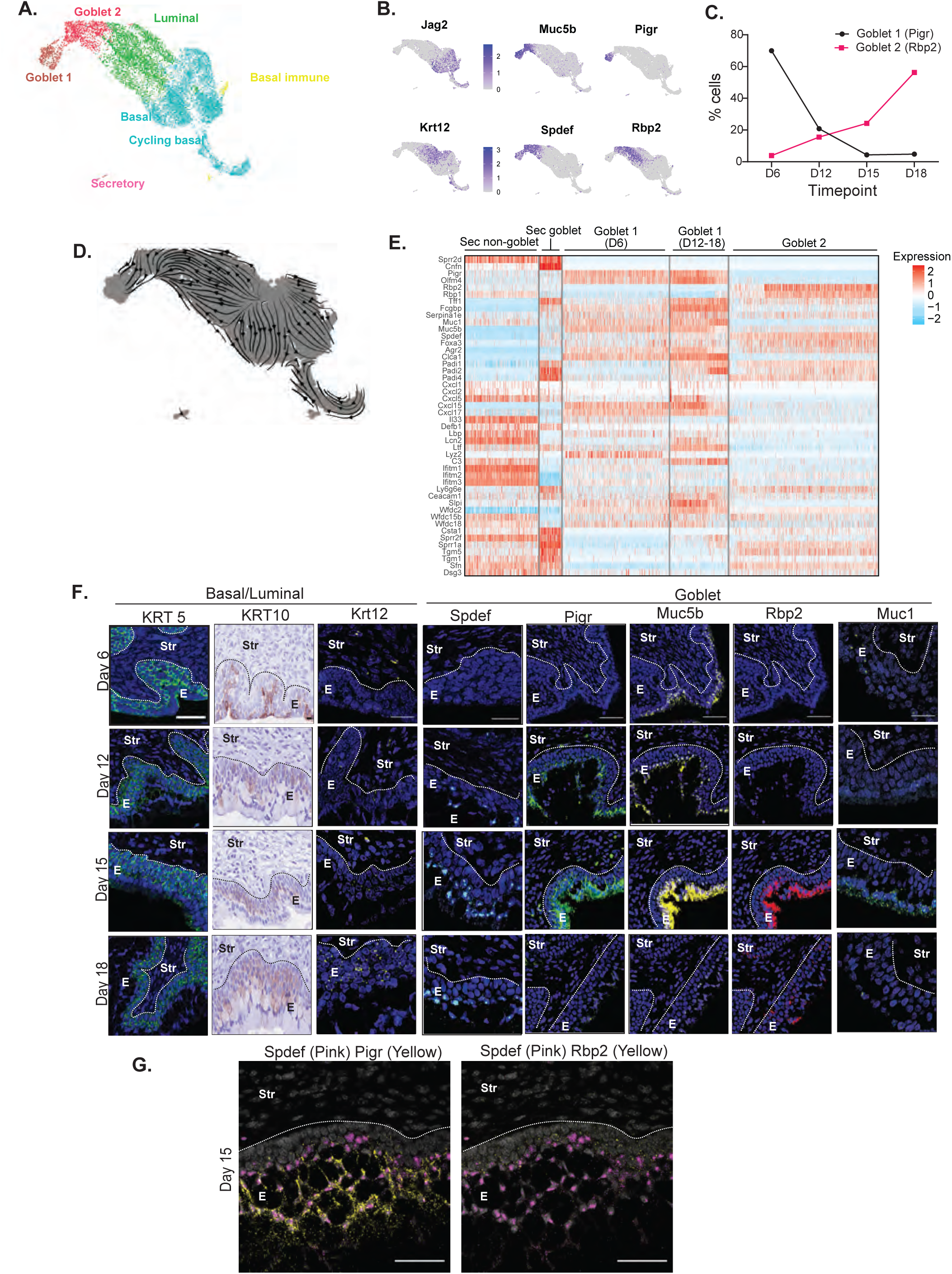
Distinct functional populations of secretory cells during pregnancy. A. UMAP visualization of epithelial cells from GD12, GD15, and GD18. B. Feature plots show the markers used to identify the different epithelia clusters. C. Abundance of goblet subtypes shift between early pregnancy to late pregnancy in scRNA-Seq libraries. Pigr^+^ goblet cells are at their highest level in GD6. Rbp2^+^ goblet cells peak at GD18. D. RNA velocity of the GD12-18 clustering shows strong directionality from luminal cells to Rbp2^+^ goblet cells, but not to Pigr^+^ goblet cells. E. Heatmap highlighting changes in gene expression across the different secretory clusters identified in the NP (Sec non-goblet and Sec-goblet) and GD6-18 clustering (Goblet 1 and Goblet 2). F. Spatial analysis of cervical epithelia subtype markers at GD6-18. Basal (Krt5), luminal (Krt10, Krt12), goblet (Spdef, Muc5b, Muc1, Pigr, Rbp2). Detection of Krt5 (green) and Krt 10 (brown) protein by IF and IHC. Detection of Krt12, Muc5b (yellow); Pigr, Muc1 (green); Spdef (blue) and Rbp2 (red) by RNAscope. DAPI(blue)-nuclei. In contrast to basal/luminal markers, expression of Spdef, Pigr, Rbp2, Muc5b, Muc1 is restricted to cells close to the lumen. E: Epithelia, Str: Stroma; Scale bar 50 µm, objective lens 40x. Representative images from three independent samples per group. G. RNAscope images showing co-analysis of Spdef (pink) with Pigr or Rbp2 (yellow) mRNA in mouse endocervical epithelia from GD15. DAPI (gray) -nuclei. E:Epithelia; Str: Stroma.Scale bar 50 µm, objective lens 40x. Representative images from two independent samples per group.

To further compare the major secretory clusters, we examined the expression status of genes relevant to cervical epithelial function (**Fig 4E**). This analysis reveals temporal and functional distinctions in secretory cell populations. Notably, the two pregnancy-specific goblet populations are distinct from each other and distinct from the NP goblet and non-goblet secretory cluster. Goblet 1 uniquely expressed olfactomedin 4, Olfm4, (Li et al., 2020; Liu et al., 2010; Schuijers et al., 2014), the chemokines, Cxcl15, Cxcl17, the calcium-activated chloride channel, Clca1 and the protease inhibitors, secretory leukocyte peptidase inhibitor (Slpi), WAP four disulfide core domain 2 (Wfdc2). Goblet 2 lacks Muc 1 expression and express keratinocyte markers (Sprr2f, Sprr2a, Tgm5, Sfn). We next evaluated the temporal patterns of goblet 1 and goblet 2 cells in the endocervix (**Fig 4F**) and spatial patterns between endocervix and ectocervix (Sup Fig 5,6,7). Krt5 and Krt10/Krt12 identify basal and non-secretory luminal cells respectively. Spdef marks both goblet 1 and goblet 2, while Pigr marks goblet 1 and Rbp2 marks goblet 2. Transcripts encoding Spdef were evident in the most luminal cell layers on GD12, 15 and 18, with the greatest number of Spdef^+^ cells on GD18. Expression of Spdef on GD6 was below the detection level for RNAscope. The strong expression of Pigr and Rbp2 on GD15 prevented the identification of goblet cells expressing only Pigr or Rbp2. As seen in **Fig 4G**, numerous Spdef^+^Pigr^+^ goblet 1 cells are identified with relatively few Spdef^+^Rbp2^+^ goblet cells. A similar pattern is noted on GD18 (**Fig S8**). Muc5b was highly expressed in the luminal layers on GD12 and GD15 with lower levels on GD6 and GD18. Muc1 transcripts were evident on days GD6, GD12 and GD15. The expansion of goblet cells in pregnancy occurs to a similar extent in both the endo- and ectocervix as seen in **Supplementary Figure 7**.

### Epithelial subtypes transition during labor towards NP estrus subtypes

To further examine epithelial cell dynamics, we expanded our analysis across NP, pregnant, and IL samples. Unexpectedly, clustering analysis shows that epithelial cells from IL samples clustered with NP rather than the pregnancy time points (**Fig 5A**). Basal and luminal cells were most concordant between the NP and IL (**Fig 5B**). Secretory cells were more divergent, with one cluster specific to NP, one specific to IL, and one shared. Among pregnancy time points, the goblet 1 cells clustered distinctly from the other time points and other clusters. While basal/luminal clusters had common markers (e.g. Jag2, Trp63) in all the time points both secretory (Muc5b, Muc1) and non-secretory (Dsg1a, Krt12) clusters were distinct between NP/IL and pregnancy time points GD 6/12/15/18 (**Fig 5C**). Analysis of keratin expression patterns (intermediate filament proteins with epithelial subtype specificity) further highlights the temporal shifts in subtype specificity from NP to pregnancy to IL (**Fig 5D**). Keratin expression and Gene Ontology pathways suggest that GD18 cells transitioning to IL rapidly adopt an NP-like state (**Fig 5D-F**). The expression of secretory markers and mucin genes (**Fig 5E**) also highlights the temporal change in secretory cell populations and the type of mucins in cervical mucus. An unappreciated diversity of antimicrobial proteins (Defensin B1 and small proline rich proteins 1 and 2) and protease inhibitors (Slpi, Wfdc2, Wfdc15b, Wfdc18, Spink 5, Spink 12) were identified with distinct temporal and subtype expression patterns (**Supp Fig 11**). Spatial analysis identifies transcriptional programs required for epithelial subtypes in the NP estrus induced in labor. RNA (Dsg1a, Spdef) and protein (Ifitm1) analysis demonstrate increased Dsg1a^+^ cells IL relative to D18 in the endo and ectocervix (**Fig 5G** **and Fig S6**). Dsg1a marks a luminal population of cells that are abundant in the estrus phase of the NP cycle. Protein staining for Ifitm1 is evident in the IL secretory cells though at reduced levels compared to the NP diestrus secretory phase. Consistent with the reduction of goblet cell differentiation, Spdef transcripts are downregulated in the IL time point relative to D18.

**Figure 5:**
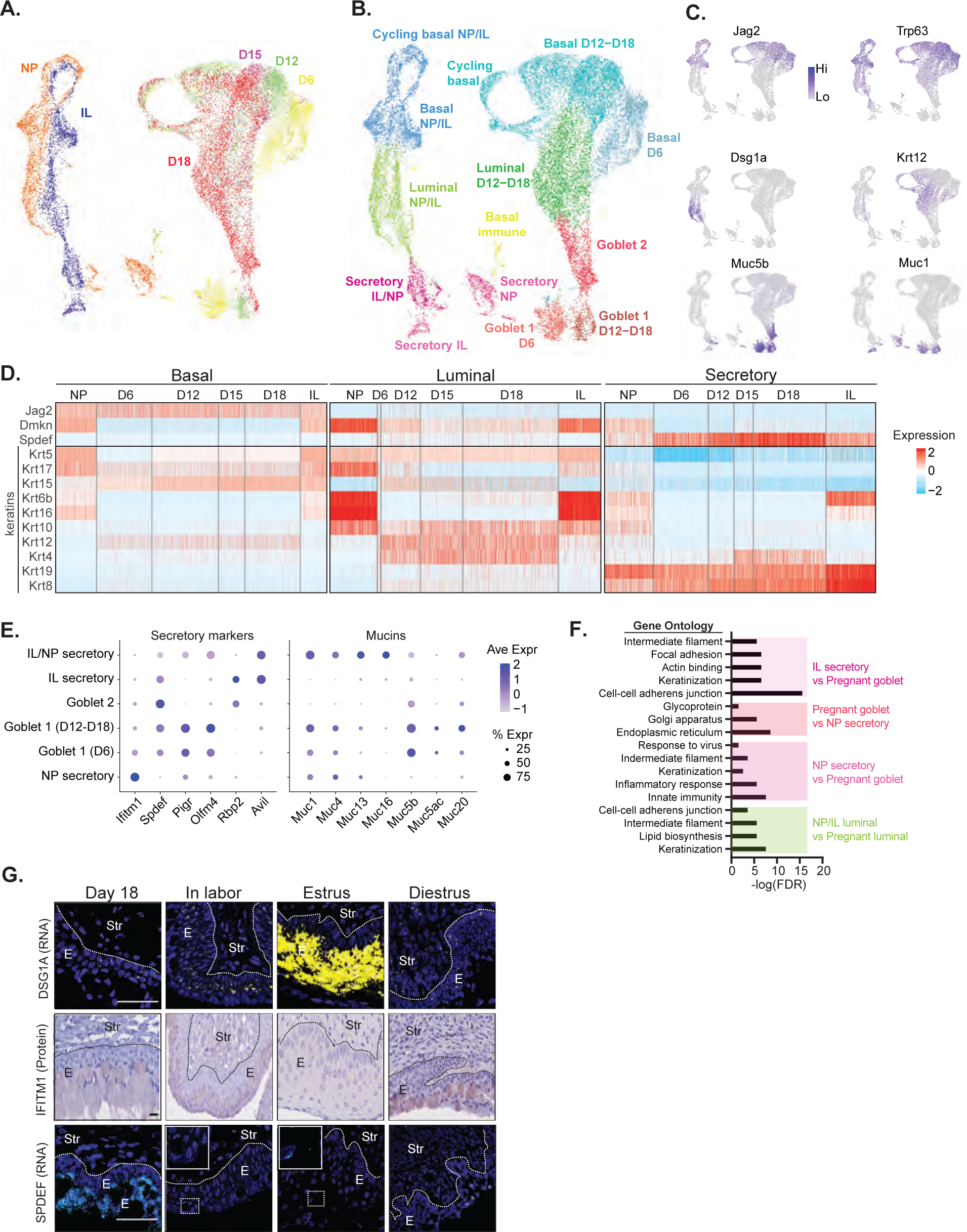
A transition of epithelial subtypes during labor towards NP estrus subtypes. A. UMAP visualization of epithelia from all time points. Cells are colored by time point. B. As in (A), but cells are colored by epithelial type. C. Feature plots showing the expression of genes that identify epithelial subtypes. D. Heatmap highlighting temporal and subtype-specific changes in keratin gene expression from basal to luminal to secretory populations across pregnancy. E. Dotplots showing the expression of secretory markers and mucins in the different secretory clusters. F. Gene Ontology analysis highlights the functional changes in the luminal and secretory populations. G. Detection of Dsg1a (yellow) and Spdef (blue) mRNA expression by RNAscope and Ifitm1(brown) protein expression by IHC on GD18, IL, and NP (estrus and diestrus). DAPI(blue) - nuclei. Inserts show a magnified view of selected areas. E: Epithelia; Str: Stroma.Scale bar 50 µm, objective lens 40x. Representative images from two independent samples per group.

### Altered cell transition states characterize epithelial barrier disruption

Our analysis identifies a pregnancy-specific expansion of luminal epithelial subtypes, including two distinct goblet populations. Next, we use these findings as a reference to define dysregulation of epithelial barrier function in mice lacking the glycosaminoglycan, hyaluronan (HA). Targeted loss of three genes (Has1, Has2, and Has3) encoding hyaluronan synthase (HAKO) in the cervix results in a morphologically disrupted secretory epithelium (**Fig 6A**), increased susceptibility to ascending infection, and premature birth (Akgul et al., 2014). To examine how HA disruption alters epithelial cell state, we performed scRNA-Seq on HAKO mice at GD15 and GD18, when cervical HA synthesis is greatest (Akgul et al., 2014; Straach et al., 2005).

**Figure 6:**
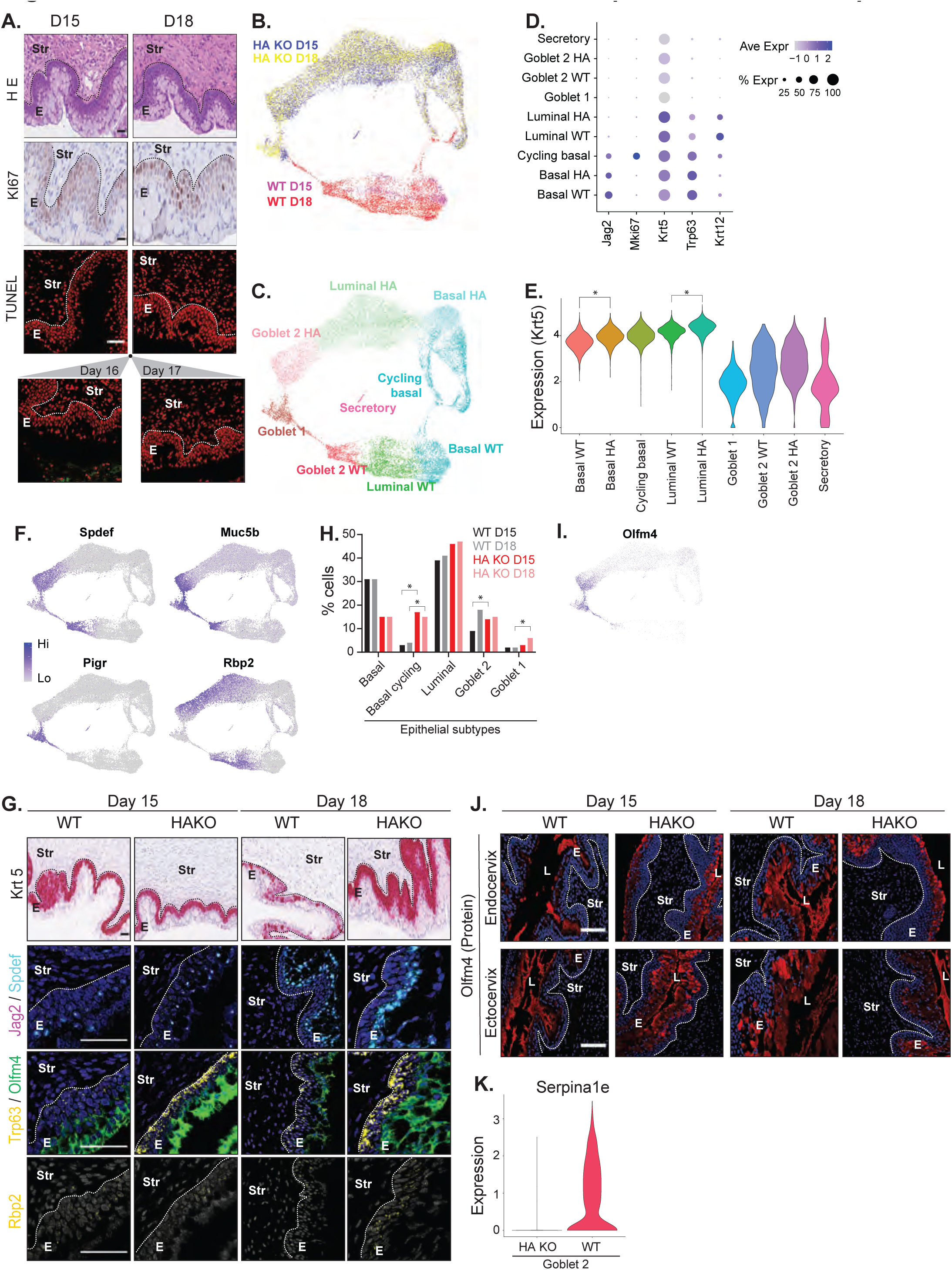
Disruptions in cell transition states characterize epithelial barrier disruption. A. Morphological evidence of cervical epithelia perturbation in HAKO on GD15 and GD18. Shown are H&E staining (top), Ki67 immunostaining for proliferation (middle), and TUNEL staining (GD15, 16, 17 and 18) for cell death (bottom) (red: nuclei, green: TUNEL^+^). E: Epithelia; Str: Stroma. Scale bar 50 µm, Representative images from three independent samples per group. B. UMAP visualization of WT and HAKO mice at GD15 and 18. Cells are colored by genotype and time point. C. As in (B), but cells are colored by epithelial type. D. Dot plot showing the expression of basal and luminal markers in WT and HAKO cells. Cycling basal, Goblet 1 and secretory subtypes are not distinguished by genotype. E. Violin plots for Krt5 expression. (* indicates p < 0.01) F. Feature plots showing the expression of goblet markers. G. Comparison of the spatial and temporal pattern of epithelial cell markers in WT and HAKO at GD15 and GD18. Basal and luminal (Krt5, Jag2, Trp63), or goblet (Spdef, Olfm4, Rbp2) transcripts were analyzed by chromogenic or fluorescent RNAScope. Panel one: Krt5 (red). Panel two: Jag2 (pink) and Spdef (blue). Panel three: Trp63 (yellow) and Olfm4 (green). Panel four: Rbp2 (yellow). DAPI stains nuclei (blue or gray). E:Epithelia; Str: Stroma. Scale bar 50 µm, objective lens 40x. Representative images from three independent samples per each group. H. The abundance of epithelial subtypes in WT and HAKO mice at GD15-18. (Hypergeometric test p-values: p_BasalCycling(D15 vs WT)_ = 9.48E-121, p_Goblet Rbb2 (D15 vs WT)_ = 7.41E–18, p_BasalCycling(D18 vs WT)_ = 5.83E-247, p_Goblet pigr (D18 vs WT)_ = 9.63E–74, * indicates p < 0.01.) I. Feature plot of Olfm4 expression. J. Olfm4 protein (red) expression in the HAKO cervical epithelia on GD15 and GD18 compared to wild-type by IF staining. DAPI stains nuclei (blue). E:Epithelia; Str: Stroma.Scale bar 50 µm, objective lens 20x. Representative images from three independent samples per each group. K. Violin plot showing loss of Serpina1e expression in goblet 2 cluster in HAKO cells.

**Figure 7:**
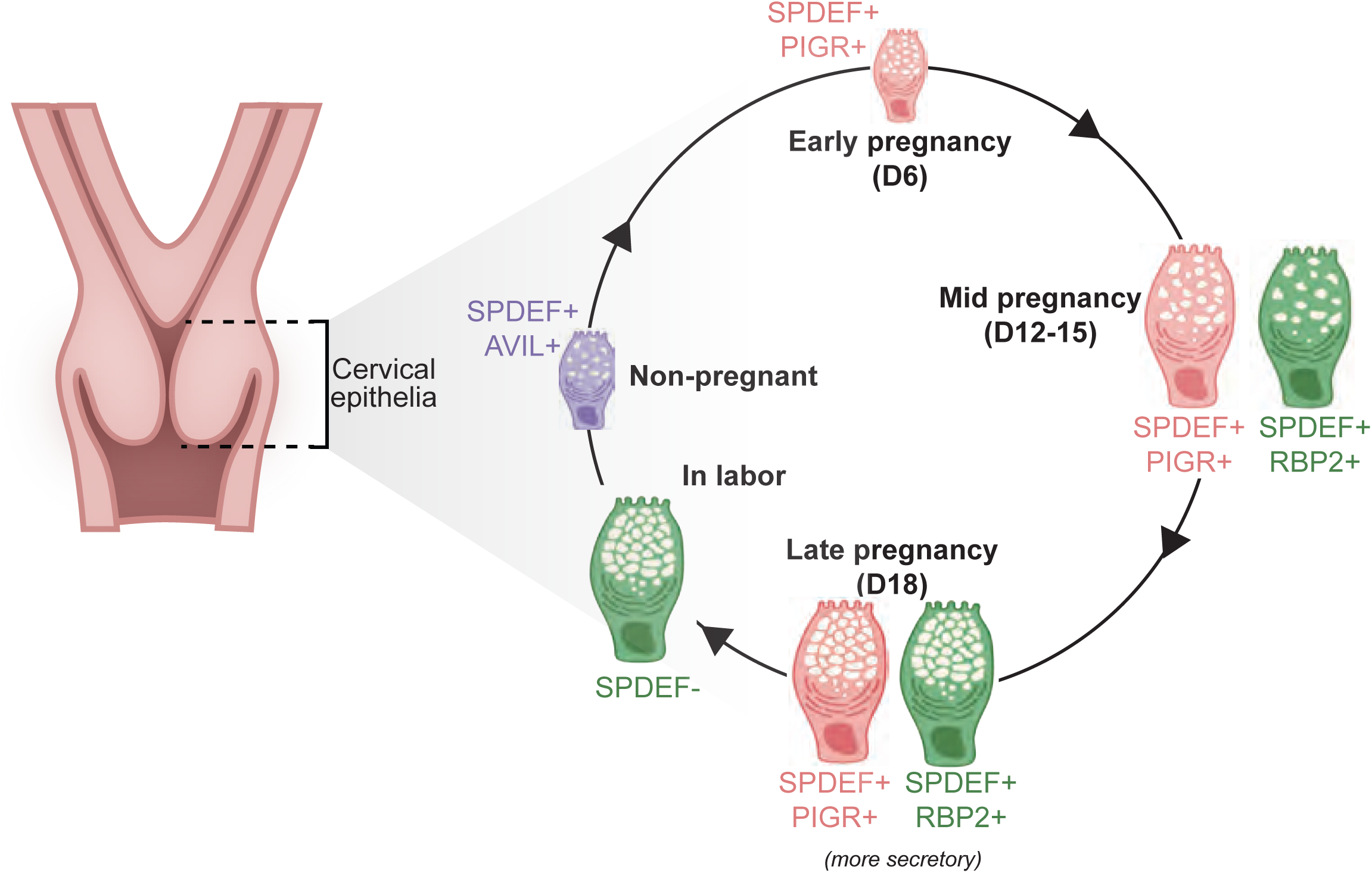
Functionally diverse goblet cell populations in mouse cervix during non-pregnant, pregnancy, and in labor. Transcriptionally, non-pregnant goblet cells are distinct from the two goblet populations identified in pregnancy. Goblet cells in NP diestrus express the marker Avil. In early pregnancy (D6), goblet cells are Pigr+ and lack Avil expression. Two distinct goblet populations are identified on gestation days 12 and 15. One is Pigr+ and the other is Rbp2+. Both populations remain on GD18 with maximal Spdef expression. Consistent with terminal differentiation of goblet cells in labor, transcriptional programs that drive goblet cell differentiation are lost and transcriptional programs in progenitor cells revert back to NP cell subtypes.

We observed that most cervical epithelial cells from HAKO mice clustered distinctly from WT mice (**Fig 6B**), with the exception of goblet cluster 1 (**Fig 6C**). Next, we examined the molecular differences between the WT and HAKO epithelial subtypes. We did not observe dramatic differences in the percentage of cells expressing characteristic marker genes for basal, luminal and goblet cells (**Fig 6D-F**). In contrast, a small increase in the average level of Krt5 expression was noted in the HAKO basal and luminal clusters relative to WT (**Fig 6D-E**) and this was confirmed in the GD18 HAKO endocervix by Krt5 spatial transcript analysis (**Fig 6G**). That loss of HA may disrupt basal cell transition cues is also suggested by the increased expression of Trp63, a transcription factor that drives differentiation of progenitor cells (Koster et al., 2004), in GD15 and 18 of HAKO relative to WT (**Fig 6G**, endocervix) and the significantly increased number of cycling basal cells relative to WT (**Fig 6H**). Immunofluorescent staining confirms increased Ki67 in the basal layer on GD15 and GD18 in HAKO relative to WT (**Fig 6A** **vs Fig1B** and **Fig S9**). These patterns were noted in both the endocervix (**Fig 6**) and the ectocervix (data not shown).

The altered cell transition states in basal cells appear to correlate with a higher number of cells marked by Olfm4 expression (**Fig 6I**). Strikingly, HAKO cells also express Olfm4 at a low level in basal and non-secretory luminal cells and at a high level in goblet 1 and goblet 2 cells, while WT mice primarily express Olfm4 in goblet 1 cells. Consistent with the single cell data, Rbp2^+^ goblet 2 cells express Olfm4 transcripts in HAKO cervices on GD15 and GD18 (**Fig 6G**), while cells co-expressing Rbp2 and Olfm4 are not visible in WT at either time point. In addition, transcripts for Olfm4 appear to increase in the GD15 and GD18 HAKO relative to WT. Olfm4 protein expression was assessed by IF (**Fig 6J**). Interestingly, the overall level of Olfm4 protein appeared similar between HAKO and WT on GD15 and GD18, yet Olfm4 was secreted into the cervical lumen in WT but remained primarily intracellular in the HAKO (**Fig 6J** **and Fig S10**). Secretion of Olfm4 has previously been demonstrated to downregulate inflammatory and immune responses in the gastrointestinal tract epithelia to limit damage to host tissue (Liu et al., 2010). Goblet 2 cell dysfunction in HAKO mice was further supported by a dramatic loss of the Serpina1e (**Fig 6K**), a serine protease inhibitor that limits protease activity and inflammatory responses (Borel et al., 2018; Lomas and Silverman, 2001). The total number of goblet cells appears similar between WT and HAKO as demonstrated by Spdef expression (**Fig 6G**). Collectively these results demonstrate hyaluronan regulates the balance between cell proliferation and subtype differentiation necessary for an immunoprotective barrier in pregnancy.

## DISCUSSION

A spatiotemporal profile of epithelial subtypes in the cervix of nonpregnant, pregnant and laboring mice provides a necessary reference from which to define molecular and cellular regulation of epithelial barrier and immune protection. Through single-cell genomic analysis, we identify pregnancy-specific epithelial subtypes with distinct temporal abundance during cervical softening and ripening in the mouse endo- and ectocervix. Complementing the single cell data, spatial analysis identifies a uniform expansion of these subtypes throughout the mouse endo and ectocervix. Despite differences between human and mouse endocervical epithelial composition, we identified numerous similarities in the expression of mucins and secreted factors as highlighted below. These findings will improve understanding of epithelial functions necessary to maintain pregnancy as well as provide a comparison to define dysregulation leading to cervical epithelial barrier disruption. This knowledge will improve understanding of host-pathogen interactions that support or compromise the function of the lower reproductive tract. Importantly, these findings will help uncover host response defects that contribute to the risk of ascending infection mediated preterm birth.

Numerous advances arise from this work. Adding to the prior understanding that the cervical epithelium has a high turnover rate in the nonpregnant cycle and is highly proliferative in pregnancy (Ramos et al., 2002), we identify shifts in the pool of proliferating epithelial cells. On gestation day 6 and to a lesser extent day 12, both basal and luminal cells are proliferative yet on GD15, 18 and IL basal cells are the dominant proliferative pool. Spatial analysis identifies these proliferating cells in both the endo- and ectocervix of the multi-layered squamous epithelia (**Fig S1**). The non-basal Ki67^+^ epithelial cells on GD6 were luminal (Krt12^+^) and goblet (Spdef^+^) subtypes. The goblet cells express markers of columnar epithelia (Krt8, Krt19) (Chumduri et al., 2021) as well as Olfm4, a marker of intestinal stem/progenitor cells and a modulator of innate immunity (Itzkovitz et al., 2011; Liu et al., 2010; Schuijers et al., 2014).

The marked shift in epithelial transcriptional programs and the pool of proliferating cells in early pregnancy resulted in distinct non-secretory and secretory luminal populations on GD6 (**Fig 3J,K**). While both non-goblet (Ifitm1^+^) and goblet (Muc5b^+^) secretory cells were identified in the NP secretory phase (diestrus), only goblet (Muc5b^+^) cells were identified on GD6. Distinct from goblet cells in the NP cervix, the pregnancy goblet 1 cluster expresses a sodium/calcium transporter (Slc12a1) and calcium channel regulator (Clca1) necessary to maintain ionic balance for mucus secretion (Erickson et al., 2016; Gagnon and Delpire, 2013; Liu and Shi, 2019). In addition, fortification of immune defense is suggested by the increased expression of protease inhibitors belonging to the whey acidic protein family (Slpi, Wfdc2), polymeric Ig receptor (Pigr) necessary for the transport of IgA antibodies into the cervical lumen (Wira et al., 2005) and Olfm4, a secreted glycoprotein with a role in innate immunity (Liu et al., 2010, 2013). Proteomic studies of cervical mucus from NP rhesus macaque (Han et al., 2021) and proteomic or limited single cell analysis of cervical samples from pregnant women highlight species conservation of noted factors observed in the goblet 1 cluster (Barnum et al., 2022; Koh et al., 2019; Kutteh and Franklin, 2001; Lee et al., 2011; Orfanelli et al., 2014).

A comparison of epithelial subtypes at four gestational time points identified two distinct goblet cell populations, designated as goblet 1 and goblet 2 (**Fig. 7**). Both computational assessments of the single cell data and spatial analysis indicate a greater abundance of goblet 1 cells from GDs 6-15 while goblet 2 cells are greatest on GDs15-18. The noted decline in goblet 1 cells on GD18 is preceded by a spike in cell death of epithelial layers that line the lumen on GDs 16 and 17 (**Fig1B**). Thus the loss of goblet 1 cells is likely achieved by cell death. Though Pigr and Rbp2 clearly mark distinct clusters in the scRNA data, spatial analysis indicates most goblet cells (Spdef^+^) have overlapping Pigr and Rbp2 expression with very few cells expressing only Pigr or Rbp2 (**Fig 4G** **and Sup** **Fig 4**). It is likely not all of the terminally differentiated goblet cells were captured by scRNA-Seq. Interestingly, while both goblet 1 and goblet 2 cells express canonical goblet cell markers (Spdef, Foxa3, Agr2) (Chen et al., 2009; Rajavelu et al., 2015), the expression of numerous genes was lower in goblet 2 (e.g. Muc1, Clca1, Cxcl15, Ltf, Slpi). In addition, goblet 2 cells express genes typically associated with luminal keratinocytes (Sprr1a, Tgm5, Sln) similar to the NP secretory clusters. Recent studies describe a bactericidal role of the small proline-rich protein family members (Hu et al., 2021; Zhang et al., 2022). Thus Sprr1a may provide antimicrobial protection in goblet 2 and NP secretory cells. Further investigation is warranted. The RNA velocity analysis (**Fig 4D**), suggests goblet cluster 2 cells (but not goblet 1) arise from the differentiation of non-secretory luminal clusters. Further studies to confirm the lineage and distinct functions of the heterogeneous population of pregnancy-distinct goblet cells in the mouse cervix are warranted. Goblet cell functional heterogeneity has been described in intestinal crypts (Nyström et al., 2021). Collectively, these studies identify the expansion of two goblet subtypes in the mouse endo- and ectocervix that synthesize a distinct mucosal network, antimicrobials (eg beta defensin and Sprr1, Sprr2) and protease inhibitors (eg Slpi, Wfdc2, Wfdc15, Wfdc18). This customized arsenal of factors in pregnancy, we suggest ensures epithelial homeostasis and provide innate immunity. In addition, the expression of immune cell chemoattractant cytokines such as Cxcl1, Cxcl2 and Cxcl5 is low or absent in goblet clusters 1 and 2 in contrast to secretory cells in the NP cervix. The noted transcriptional changes would provide mucosal immunoprotection without eliciting proinflammatory responses that could induce preterm birth.

Goblet cell diversity with distinct antimicrobial molecule secretion, and mucus gene expression may collectively contribute to ensuring barrier function during pregnancy. This process of inflammatory remodeling is potentially similar to that described in the barrier epithelium of the intestine and airways (Medzhitov, 2021). Importantly, the noted ability of the pregnancy-specific goblet subtypes to regulate the mucus structure, and display differential expression of chemokines/cytokines, antimicrobials, protease inhibitors, complement and immunoglobulin transporters motivates further studies to define their regulation and function in the physiology of term pregnancy and in spontaneous preterm birth (sPTB) in women. In further support, numerous studies highlight the association of sPTB in women with a diverse vaginal microbiota, reduced *Lactobacillus* content and a maternal host immune response characterized by increases in proinflammatory cytokines, chemokines, antimicrobials, IgM/IgG and complement activation (Chan et al., 2022; Elovitz et al., 2019; Flaviani et al., 2021; Florova et al., 2021). In addition, reduced mucus permeability characterizes cervical mucus of women who deliver preterm relative to term (Critchfield et al., 2013).

The marked shift in the transcriptional program of epithelial cells during labor emphasizes the dynamic nature of epithelial cell homeostasis. Our analysis of epithelial cells from NP, pregnant and IL, clearly demonstrate a marked change in the transcriptional program of epithelial cells during labor to regenerate epithelial subtypes present in the NP cervix. In support, spatial studies demonstrate increased transcription of Dsga1, a marker of luminal keratinocyte cells (Harmon et al., 2013) present in the NP estrus stage of the cycle. The dynamic temporal change in epithelial subtypes from NP to pregnancy and the return to an NP pattern in labor is reflected in the change in expression of keratin intermediate filament proteins (Harmon et al., 2013; Karantza, 2011) as seen in the heatmap in **Fig 5D**. As noted earlier, the secretory populations in all libraries express the columnar cell marker Krt8. In addition, these secretory clusters were noted to express the squamous marker Krt5. Confirmatory spatial studies identify Krt5 protein in all layers of the epithelia through the mouse endo- and ectocervix (**Supp Fig 2 and 3**). These findings raise the possibility of a cellular plasticity by which squamous cells transdifferentiate into the Krt8+ secretory cells. Future lineage tracing studies will distinguish between a “physiological metaplasia” whereby squamous progenitor cells transform into secretory columnar cells or the alternative possibility that multiple progenitors give rise to diverse epithelial subtypes identified during pregnancy. The ability of a plastic bi-potent cervical epithelial stem/progenitor cell to give rise to both squamous and columnar epithelia in humans and mice continues to be debated as demonstrated in recent studies which support(Zhao et al., 2021) and refute(Chumduri et al., 2021) this hypothesis.

Future studies to define steroid hormone regulation of epithelial subtype diversity are needed. The role of progesterone and estrogen via activation of their nuclear receptors, progesterone receptor (Pgr), estrogen receptor alpha and beta (Erα, Erβ) in the regulation of nonpregnant cervical epithelial proliferation, differentiation, mucosal composition and function is well documented (Galand et al., 1971; Gipson et al., 1997; Ludwig, 1989; Ramos et al., 2002; Wang et al., 2001; Wira et al., 2015). Goblet cells are demonstrated to expand in the NP-proestrus cycle in mice with a Muc5b-GFP reporter thus indicating P4 and E2’s ability to regulate specific epithelial subtypes (Portal et al., 2017). Ongoing studies will leverage scATAC-Seq data to identify epithelial subtype-specific direct and indirect transcriptional targets of P4 and E2 through pregnancy and labor. However, one drawback is that the scATAC-Seq data presented here lacks sufficient coverage of all epithelial subtypes sampled by scRNA-Seq, likely due to differences between the cellular isolation protocol of scRNA-Seq and nuclear isolation protocol of scATAC-Seq.

Using mice lacking cervical hyaluronan (HAKO), we identify an untimely proliferation of basal cells with enhanced expression of the transcription factor Trp63, essential for the proliferative potential of stem/progenitor stratified epithelia (Senoo et al., 2007). Notably, these cells differentiate into goblet cells expressing canonical markers (eg Spdef, Muc5b, Foxa3, Agr2), though goblet cell organization and some aspects of function (e.g. loss of Serpina1e transcripts and luminal Olfm4 secretion) are perturbed. While further interrogation of our data is required to define the hyaluronan signaling cues that are driving timely proliferation, differentiation and organization of the mucosal epithelia, these findings highlight the necessity of finely-regulated temporal changes in epithelial subtypes required for the maintenance of an optimal mucosal barrier through pregnancy.

The advent of single cell technology has broadened our understanding of epithelial cell diversity necessary to maintain tissue homeostasis and the potential of host-microbe interactions to modify epithelial subtype populations or function in numerous tissues (Bomidi et al., 2021; Parikh et al., 2019). Our studies in the mouse cervix not only identify subtype heterogeneity but also suggest epithelial cell remodeling is necessary to maintain a dynamically shifting state of homeostasis in pregnancy and labor. The findings of the current study provide a framework to determine if perturbations in finely regulated signaling cues that drive epithelial proliferation, differentiation and organization contribute to the risk of ascending infection. This research may lead to the identification of biomarkers that identify loss of cervical epithelial cell health in pregnancy and the development of preventive therapies that will reduce the risk of ascending inflammation-mediated preterm birth.

## ACKNOWLEDGEMENTS

We acknowledge the BioHPC computational infrastructure at UT Southwestern for providing high performance computing and storage resources that have contributed to the research results reported within this paper. We would like to acknowledge the assistance of the UT Southwestern Histo-Pathology Core and Quantitative Light Microscopy Core, a Shared Resource of the Harold C. Simmons Cancer Center, supported in part by an NCI Cancer Center Support Grant, 1P30 CA142543-01. We thank the UT Southwestern Medical Center Whole Brain Microscopy Facility (RRID:SCR_017949) for assistance with whole slide imaging. This work is supported by the Burroughs Wellcome Fund (1019804) and 1P01HD087150-01A1 – Project 4 to M.M. G.C.H is supported by CPRIT (RP190451), NIH (DP2GM128203, UM1HG011996), the Burroughs Wellcome Fund (1019804), and the Green Center for Reproductive Biological Sciences. A.C. and S.M. share equal co-authorship, and author order was determined by tossing a Hot Potato.

## AUTHOR CONTRIBUTIONS

M.M. and G.C.H. designed the experiments. L.W. performed the genomic experiments. A.C. performed computational analysis of genomic datasets, with assistance from E.S. S.P.M. performed single cell isolation procedures, prepared sections and completed staining for RNA and protein studies and image analysis for H&E, TUNEL, IHC, IF and RNAscope. All authors prepared the manuscript. M.M. and G.C.H. secured funding to support this project and provided intellectual support for all aspects of the work.

## DATA AND CODE AVAILABILITY

The sequencing data generated in this study have been deposited to the Gene Expression Omnibus (GEO) under the accession (GSE196529, reviewer token: grqjuogovterpqf).

## METHODS

### Mice

All the experimental procedures with mice were approved by the Institutional Animal Care and Use Committee at the University of Texas Southwestern Medical Center. Mice were housed under a 12-hour light/dark cycle at 22°C in a non-barrier facility and were provided with food and water ad libitum. Mice used in this study were 2- to 6-month-old and nulliparous. *Nonpregnant* mice were assessed for stage of the estrus cycle. To determine the cycle, vaginal smears were taken and analyzed for the presence or absence of cornified epithelia, nucleated epithelia and leukocytes (Byers et al., 2012). To collect tissues from *pregnant and in labor*, females were housed with males overnight and checked for vaginal plug in the morning. The day of plug formation was counted as day 0 with birth occurring on gestation day 19. In labor (IL) samples were collected on d19 after delivery of 1-2 pups. Mice used in this study were of C57B6/129SV strain. As previously described, the Has1/2/3 knockout mouse line has global loss of hyaluronan synthase 1 and 3 (Has1^-/-^ and Has3^-/-^) and cell-specific loss of Has2 in cells expressing progesterone receptor (Has2^fl/fl^; PRCre^+/-^) (Akgul et al., 2014).

### Tissue collection

For the single cell studies cervices were collected by dissection below the transition zone (TZ) (Fig 1) and all vaginal tissue was removed. For spatial studies, cervices with vagina and uterine horns were collected for longitudinal sections with the cervical canal evident throughout each section and in the same longitudinal plane as the uterine horns.

### Single cervical cell dissociation and library preparation

Cells for the single-cell RNA libraries, were isolated from cervices of NP (diestrus (n=7) and estrus (n=2)) and gestational days 6 (n=5), 12 (n=4), 15(n=3), 18(n=3), and IL (n=3). The number of pooled cervices used for each library is indicated. Tissues were minced with a sterile razor. Minced cervices were dissociated with 10mg/ml cold protease (Sigma-Aldrich) with 1mg/ml DNase I (Roche) and 0.5 M EDTA (Fisher Scientific). After a 60 min incubation at 4°C, cells were filtered through a 40 µm cell strainer (BD Biosciences) and washed twice with fresh ice cold 1x PBS and then resuspended in 1x PBS +0.005% BSA. Cell viability and numbers were assessed by trypan blue stain and counted using Hemocytometer. scRNA-Seq was carried out as recommended by the 10X Genomics standard operating procedure.

Nuclei for the single cell ATAC-Seq (scATAC-Seq) libraries were isolated from frozen mouse cervices of NP metestrus (n=2) and gestational days 6 (n=3), 12 (n=2), 15 (n=2), 18 (n=1) and IL (n=1) following 10X Genomics protocol. Briefly, cervices were homogenized in 0.1X Lysis buffer with a pellet pestle for 5 minutes on ice. Reaction was stopped by adding the wash buffer to the lysed cells and suspensions were filtered through a 40um cell strainer (BD Biosciences). Nuclei concentration was determined by DAPI and Ethidium Homodimer-1 dyes using a Countess II FL Automated Cell Counter. scATAC-Seq was conducted as recommended by 10X Genomics User Guide.

### Spatial analysis in tissue sections

All tissues were fixed in 4% paraformaldehyde (Sigma-Aldrich) in 1x PBS for 24 hrs at 4°C then transferred to 1x PBS. Tissues were embedded in paraffin and 5 µm thick sections were cut. TUNEL staining (Promega Dead End Fluorometric TUNEL kit) and Hematoxylin (HE) staining were performed by the Histo-Pathology Core at UT Southwestern Medical Center.

#### Immunohistochemistry

Paraffin sections were deparaffinized using standard procedure. After antigen retrieval using sodium citrate buffer (10mM pH 6) for 20 mins at 4°C, sections were subjected to 0.5 % hydrogen peroxide solution at RT for 20 mins (H1009, Sigma) in methanol to quench endogenous peroxidases. Sections were blocked with 10% normal goat serum (50062Z, Life Technologies) for 30 mins at RT and rinsed in 1x PBS. Subsequently, sections were incubated with streptavidin and biotin blocking kit (SP-2002, Vector Laboratories) for 20 mins at RT. After washing with 1x PBS the sections were incubated with primary antibody overnight at 4°C. The next day, slides were washed with 1x PBS and incubated with biotinylated secondary antibody for 1 hr at RT followed by VECTASTAIN ABC-HRP reagent, peroxidase (PK-7100, Vector Laboratories) for 30 mins followed by Vector NovaRED substrate peroxidase solution (SK-4800, Vector Laboratories) at RT. After rinsing the slides with tap water sections were counterstained using Hematoxylin (72804, Epredia) for 10 sec. Slides were washed in 1x PBS and dehydrated in 70% ethanol, 100% ethanol and Xylene. Finally, sections were covered with Cytoseal (8312-4, Thermo scientific) and dried for imaging.

#### Immunofluorescence

Following deparaffinization and antigen retrieval as described above, tissue sections were blocked with 10% normal goat serum (500627, Thermo scientific) for 30 mins at RT. Sections were rinsed in 1x PBS and incubated with primary antibody overnight at 4°C. The next morning, sections were washed in 1x PBS and incubated with secondary antibody for 30 mins at room temperature. Sections were mounted with Prolong Gold anti-fade reagent with DAPI (P36935, Invitrogen) and sealed with a coverslip.

#### RNAscope

Chromogenic or Multiplex fluorescent Kit (Advanced cell diagnostics 320511 or 323100) was used as per the manufacturer’s instructions. Briefly, slides were deparaffinized in xylene, followed by rehydration in a series of ethanol washes. Following citrate buffer, and antigen retrieval, slides were treated with protease plus (Advanced Cell Diagnostics) in a HybEZ hybridization oven (Advanced Cell Diagnostics). Probes directed against genes of interest (mRNA) and control probes were applied at 40 °C in the following order: target probes, preamplifier, amplifier; and label probe for 10 min. After each hybridization step, slides were washed two times at RT. The target retrieval boiling time was adjusted to 15 mins and incubated with protease plus at 40°C for 30 mins. For chromogenic, hybridization signals were detected using diaminobenzidine (DAB). The RNA signal was identified as red punctate dots. For fluorescent, slides were mounted with DAPI and Prolong Gold antifade reagent (P36930) and sealed to dry and stored at 4°C until imagining

### Image Analysis

For Hematoxylin and immunohistochemistry, the whole slides were scanned using the Hamamatsu Nanozoomer 2.0 HT and viewed using NDP View2 software. For TUNEL, immunofluorescence and RNAscope, images of longitudinal cervical sections were taken with Zeiss LSM780 inverted confocal microscope at 20x and 40x magnifications.To evaluate the temporal changes, all the opal labelled RNAscope images were captured using the same detector gain and laser power.To ensure we could compare relative expression for all the time points, the captured images were rescaled linearly for each channel across all time points. Each individual image was processed with scale bars using Image J software.

### Antibodies and Probes

Antibodies used for immunohistochemistry and immunofluorescence: rabbit anti-Keratin 10 (1:100, Abcam ab76318), mouse anti-IFITM1 (1:50, Proteintech 60074-1), rat anti-Ki67 (1:200, ThermoFisher scientific 14-5698-82), chicken anti-Keratin 5 (1:100, Biolegend 905901), and Biotinylated goat anti-rabbit IgG (1:200, Vector laboratories BA-1000), goat anti-mouse IgG (1:200, Vector laboratories BA-9200) and goat anti-Rat IgG (1:200, Vector laboratories BA-9400), Donkey anti-chicken Alexa Fluor 488 (1:600, Jackson ImmunoResearch 703-545-155) conjugated secondary antibodies. Probes used for RNAscope (Advanced Cell Diagnostics): *Krt5* (415041), *Avil* (C1, 498531), *Dsg1a* (C2, 842861), *Spdef* (C2, PN 544421), *Muc5b* (C2, 471991), *Muc1* (C1, 421871), *Pigr* (C1, 552591), *Rbp2* (C3, 444091), *Olfm4* (C2, 311831).

### Single cell data clustering

Seurat V3.2 was used for scRNA-seq data analysis (Stuart et al., 2019). The standard workflow was used for clustering. UMAP clustering was first run on the entire dataset (each timepoint individually). Then DoubletFinder V2.0.3 was used to identify any doublets. Cells that were identified as doublets were removed. After doublet removal, the cell type of each cluster was determined. The expected cell types include epithelial, stromal, endothelial, immune, and blood cells. Using known genes, all these expected cell types were found in the single-cell data. Any clusters that were not identifiable as a particular cell type were removed from the analysis. These clusters exhibited low gene counts and were of poor quality. A 20% mitochondrial cutoff was also applied to remove any cells with a higher than expected percent of mitochondrial reads indicating cells of poor quality. After the data was cleaned up, clustering was re-run using the Seurat pipeline on each time point. The clusters identified in each time point were used for further downstream analysis. Seurat functions, FindMarkers and FindAllMarkers, were used to identify differentially expressed genes between clusters. scVelo 0.2.2 was used for RNA velocity analysis after clustering was performed.

### Single Nuclei ATAC-Seq Data Analysis

ArchR 2.0 was used to analyze the snucATAC-seq data (Granja et al., 2021). The cells from each timepoint were filtered using a TSS enrichment score cut-off above 4 and a unique nuclear fragments cut-off above 1000. Doublets were also identified and filtered using the AchR workflow. After filtering and doublet removal, each timepoint NP, day 6, 12, 15, 18, and IL had 3288, 7806, 5824, 6530, 2312, and 4998 cells respectively. Cells from all time points were clustered using UMAP, and gene scores and deviation scores were calculated using the standard ArchR workflow. Cell type identity for each cluster was determined using the gene score for known genes, and those used in the scRNA-seq data analysis.

## SUPPLEMENTAL FIGURE LEGENDS

**Figure S1.**
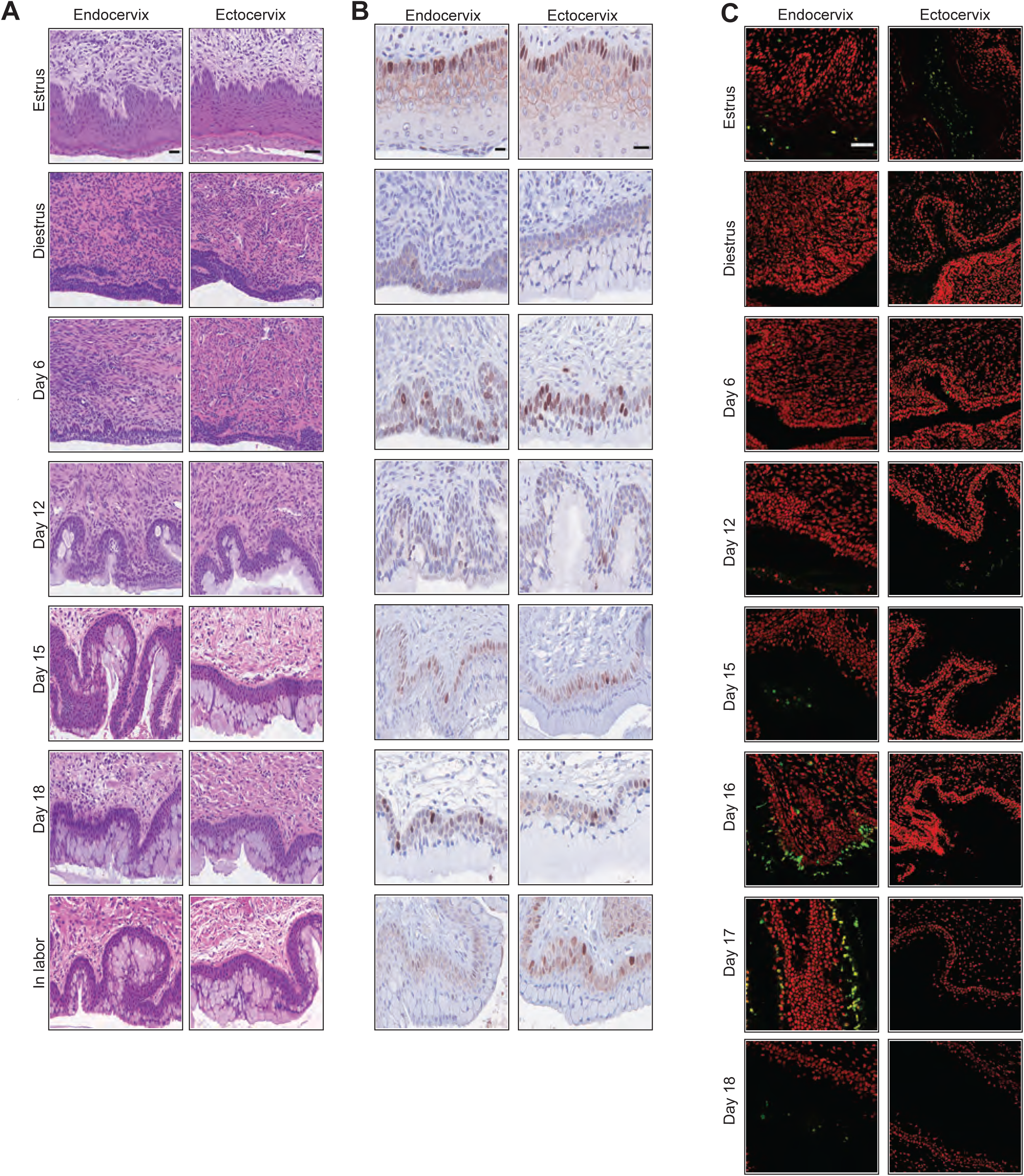
**A.** H&E histology of endo- and ecto-cervical epithelia in NP, pregnant (D6, D12, D15 and D18), and in labor (IL) mice. The representative sections are shown at 60x magnification from three independent samples per group. E: Epithelia; Str: Stroma. Scale bar 50 µm B. Immunostaining for Ki67 (brown) identify proliferating cells in endo- and ecto-cervical epithelia in NP, pregnant (D6, D12, D15 and D18), and in labor mice. The representative sections are shown at 60x magnification from three independent samples per group. E: Epithelia; Str: Stroma. Scale bar 50 µm C. TUNEL staining in endo- and ecto-cervical epithelium of NP, pregnant (D6, D12, D15, D16, D17 and D18), and in labor mice. (red: nuclei, green: TUNEL^+^). The representative sections are shown at 20x magnification from three independent samples per group. E: Epithelia; Str: Stroma. Scale bar 50 µm

**Figure S2.**
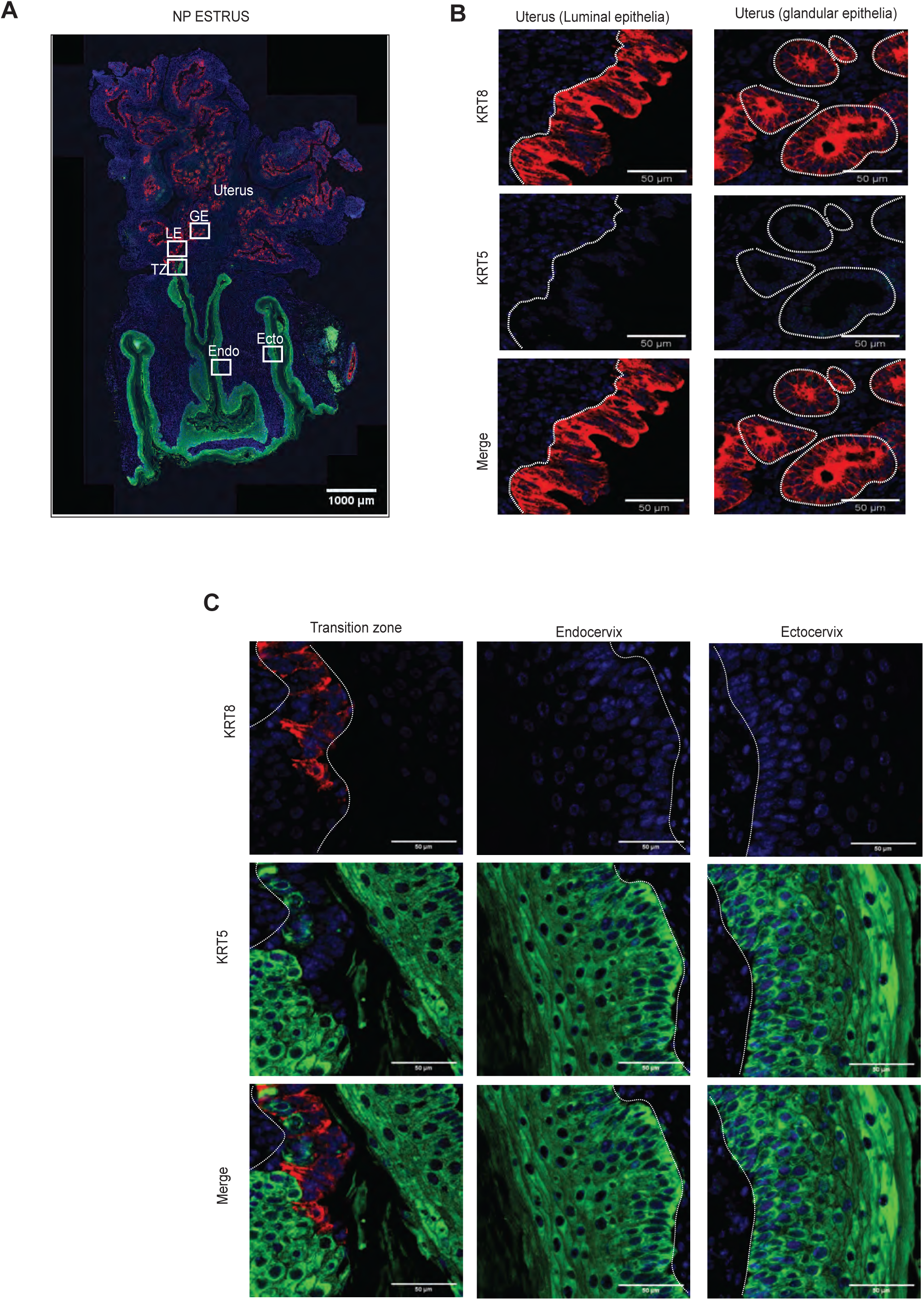
**A.** Confocal images of the mouse reproductive tract from **nonpregnant estrus** stained with columnar marker Krt8 (red) and stratified squamous marker Krt5 (green) and nuclei in blue. Boxed areas are magnified in B and C. Representative of biologically independent experiments from 3 mice.TZ:Transition zone; Endo:endocervix; Ecto:ectocervix; LE: luminal epithelium; GE: glandular epithelium. B. Shown are magnified images of the uterine luminal and glandular epithelia stained with columnar marker Krt8 (red) and stratified squamous marker Krt5 (green) and nuclei in blue. C. Magnified images of the transition zone, endocervix and ectocervix stained with columnar marker Krt8 (red) and stratified squamous marker Krt5 (green) and nuclei in blue.

**Figure S3.**
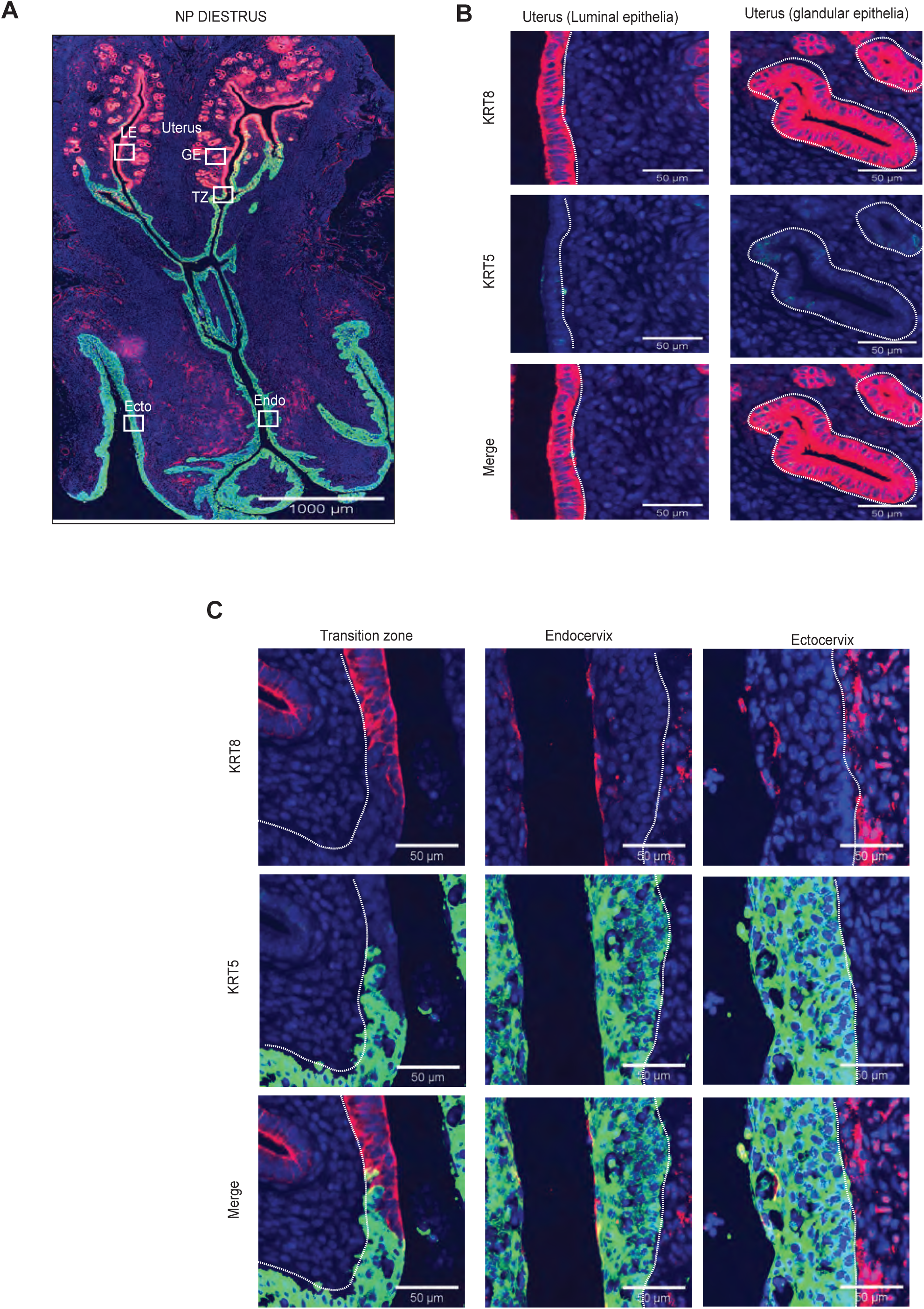
**A.** Confocal images of the mouse reproductive tract from **nonpregnant diestrus** stained with columnar marker Krt8 (red) and stratified squamous marker Krt5 (green) and nuclei in blue. Boxed areas are magnified in B and C. Representative of biologically independent experiments from 3 mice.TZ:Transition zone; Endo:endocervix; Ecto:ectocervix; LE: luminal epithelium; GE: glandular epithelium. B. Magnified images of the luminal and glandular epithelia lining the uterus stained with columnar marker Krt8 (red) and stratified squamous marker Krt5 (green) and nuclei in blue. C. Magnified images of the transition zone, endocervix, and ectocervix stained with columnar marker Krt8 (red) and stratified squamous marker Krt5 (green) and nuclei in blue.

**Figure S4.**
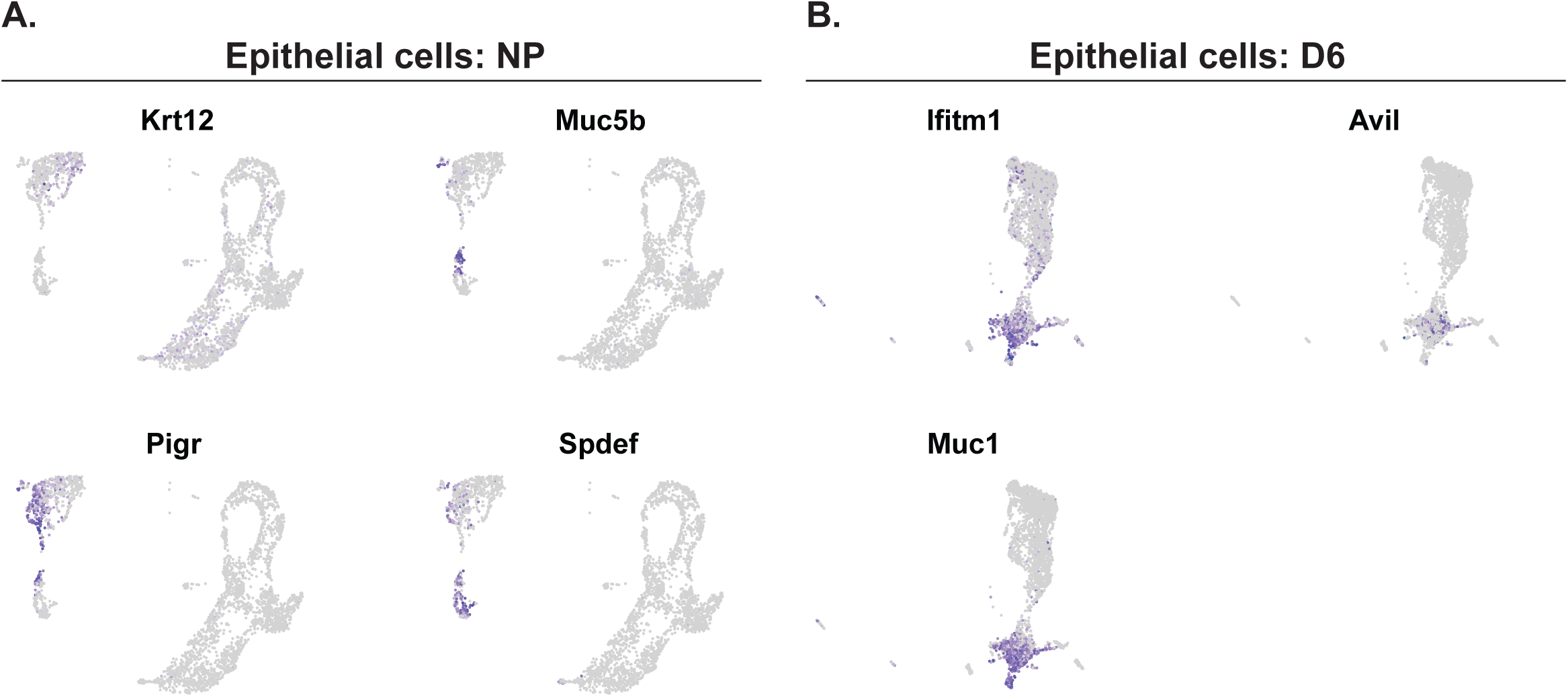
**A.** Feature plots showing the expression of luminal and goblet markers in the NP time point. B. Feature plots showing the expression of secretory markers in the D6 time point.

**Figure S5.**
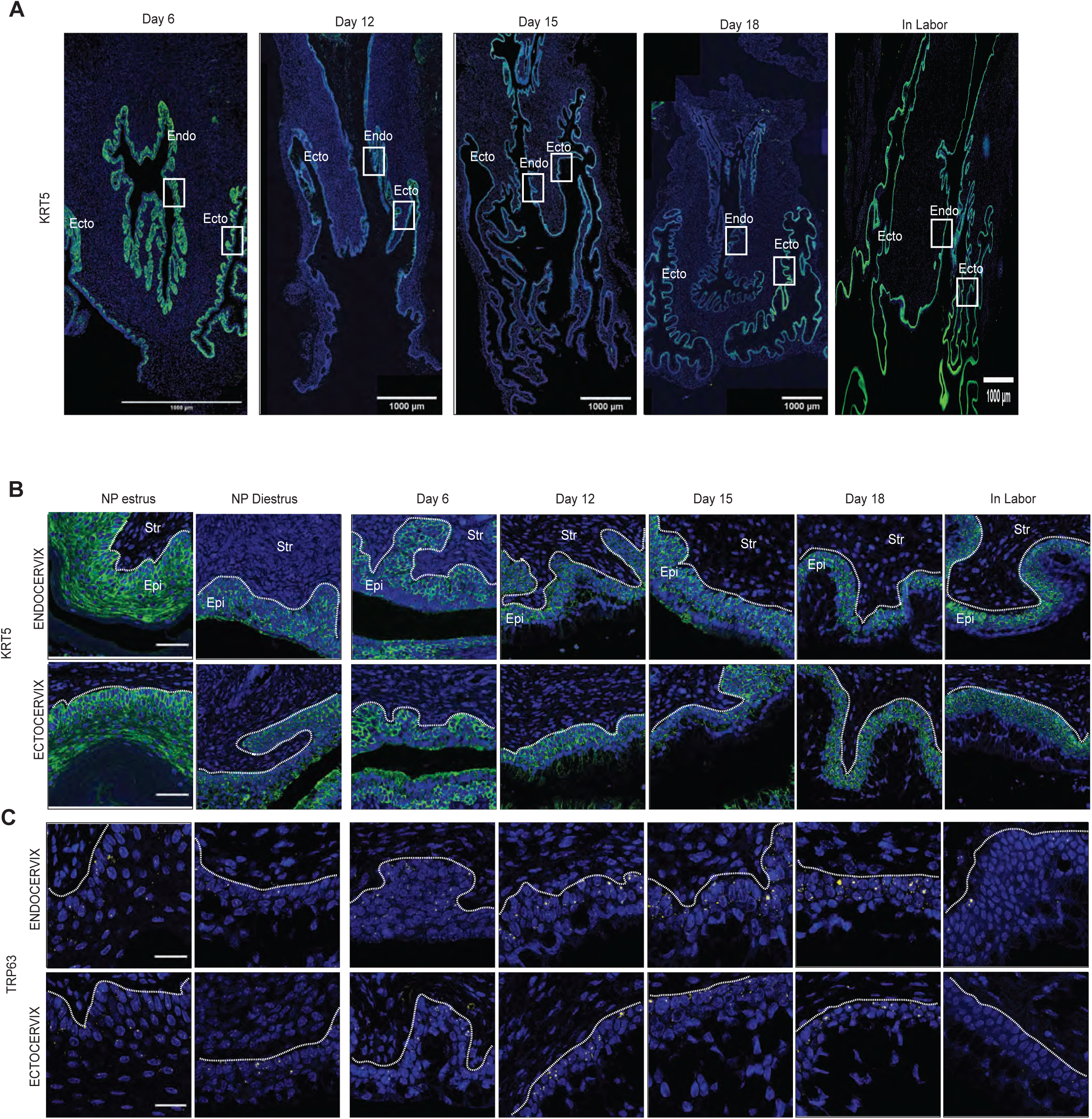
**A.** Confocal images of the mouse reproductive tract during pregnancy (D6, D12, D15 and D18), and in labor (IL) stained with stratified squamous marker Krt5 (green) and nuclei in blue. Boxed areas are magnified in Panel B. Representative of biologically independent experiments from 3 mice. Endo:endocervix; Ecto:ectocervix. B. Spatial analysis of the basal marker Krt5 (green) protein immunostaining of endo- and ectocervical epithelia in NP, pregnant (D6, D12, D15 and D18), and in labor (IL) mice.The representative sections are shown at 40x magnification. Epi: Epithelia; Str: Stroma. Scale bar 50 µm C. Spatial analysis of the transcription factor Trp63 mRNA (yellow) at endo- and ectocervical epithelia in NP, pregnant (D6, D12, D15 and D18), and in labor (IL) mice. The representative sections are shown at 40x magnification. Scale bar 50 µm

**Figure S6.**
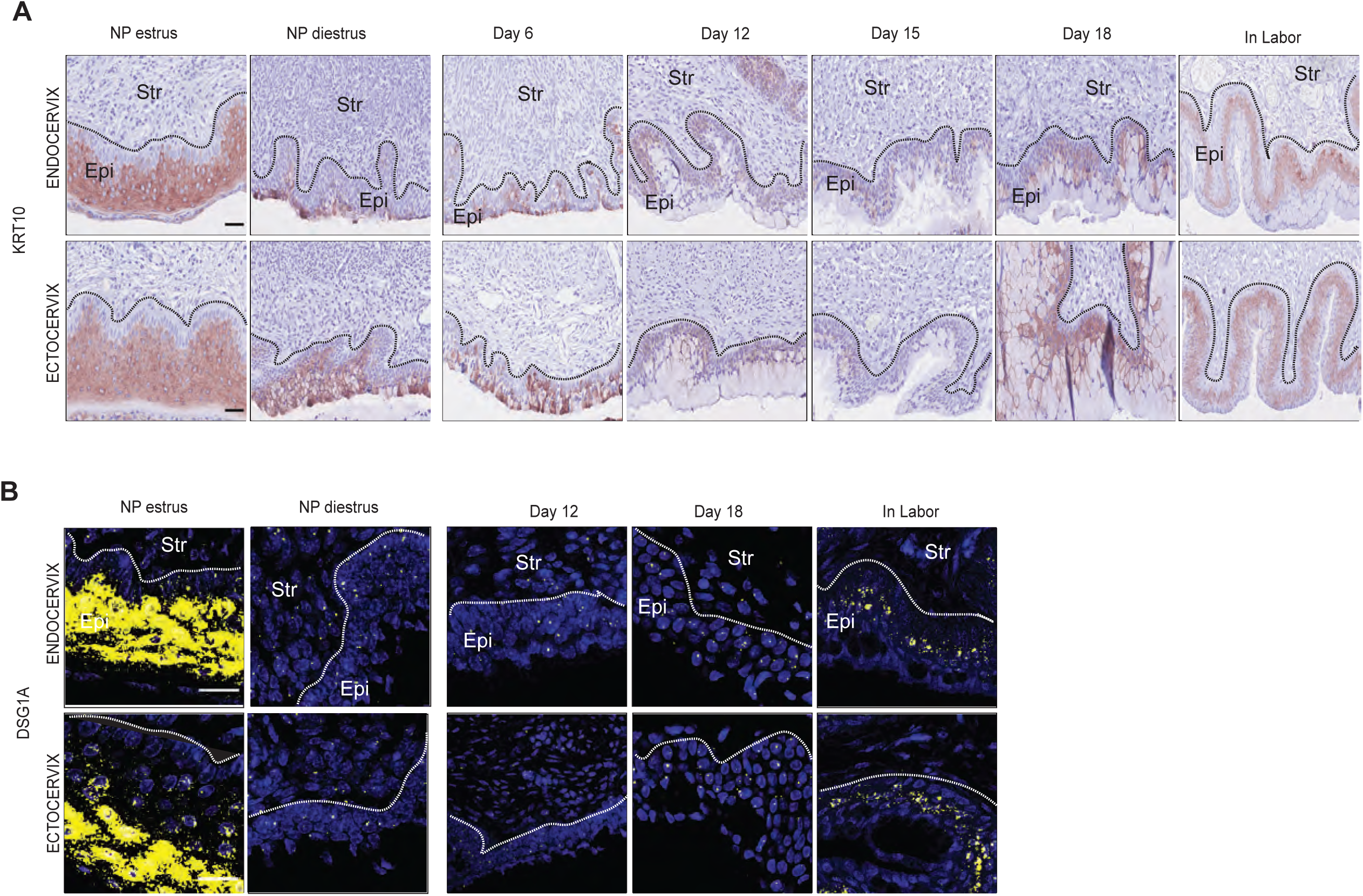
**A.** Immunostaining for luminal marker Krt10 (brown) in endo- and ectocervical epithelia in NP, pregnant (D6, D12, D15 and D18), and in labor mice. The representative sections are shown at 40x magnification from three independent samples per group. Epi: Epithelia; Str: Stroma. Scale bar 50 µm B. Spatial analysis of the luminal marker Dsg1a (yellow) mRNA expression by RNAscope in endo- and ecto-cervical epithelia in NP, pregnant (D12 and D18), and in labor (IL) mice. The representative sections are shown at 40x magnification. Epi: Epithelia; Str: Stroma. Scale bar 50 µm

**Figure S7.**
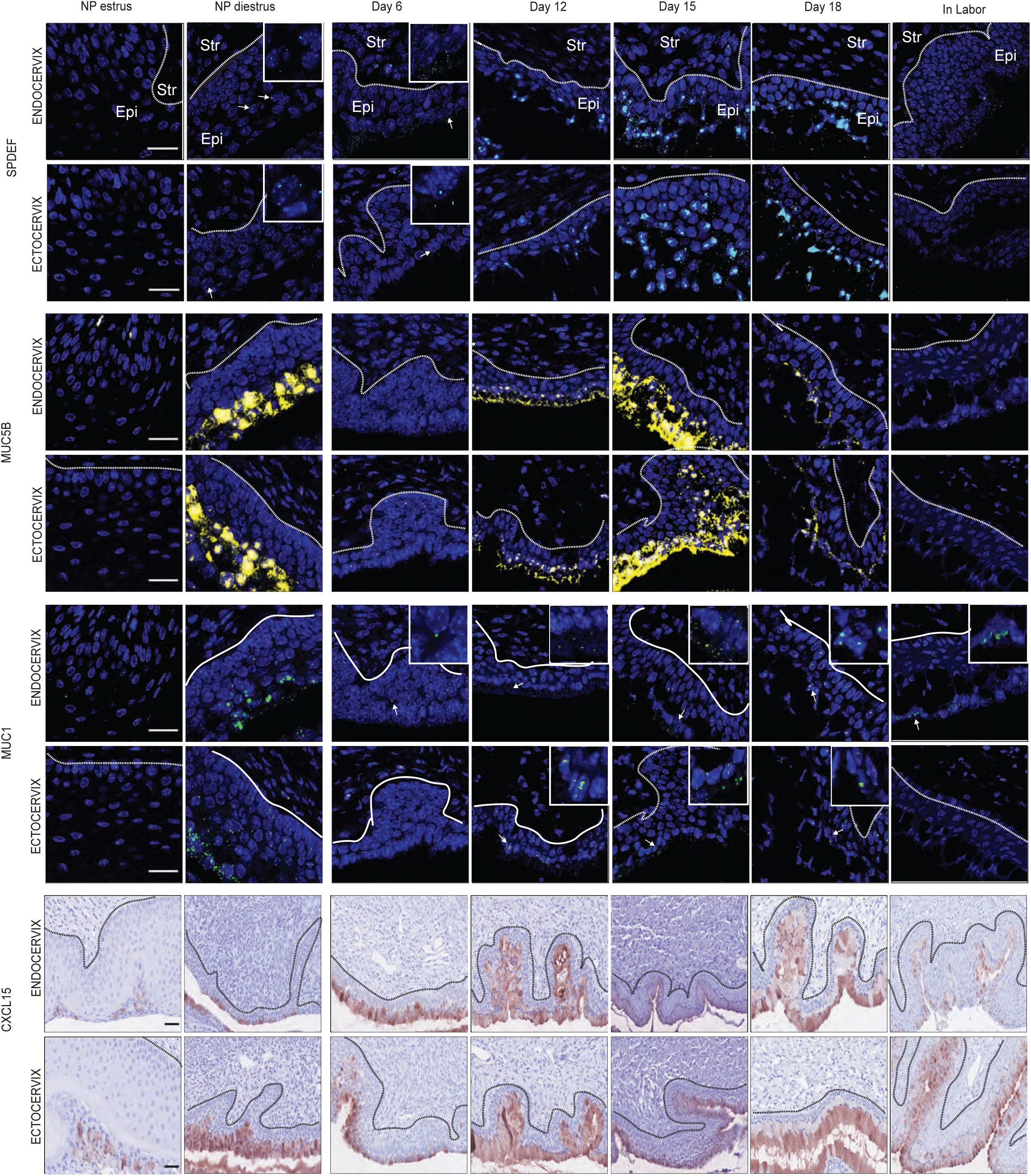
MRNA expression of secretory markers Spdef (blue), Muc5b (yellow), Muc1 (green) by RNAscope and protein expression CXCL15 (brown) by immunostaining in endo- and ectocervical epithelia in NP, pregnant (D6, D12, D15 and D18), and in labor (IL). The representative sections are shown at 40x magnification. Magnification of white arrows are shown in inserts. Epi: Epithelia; Str: Stroma. Scale bar 50 µm

**Figure S8.**
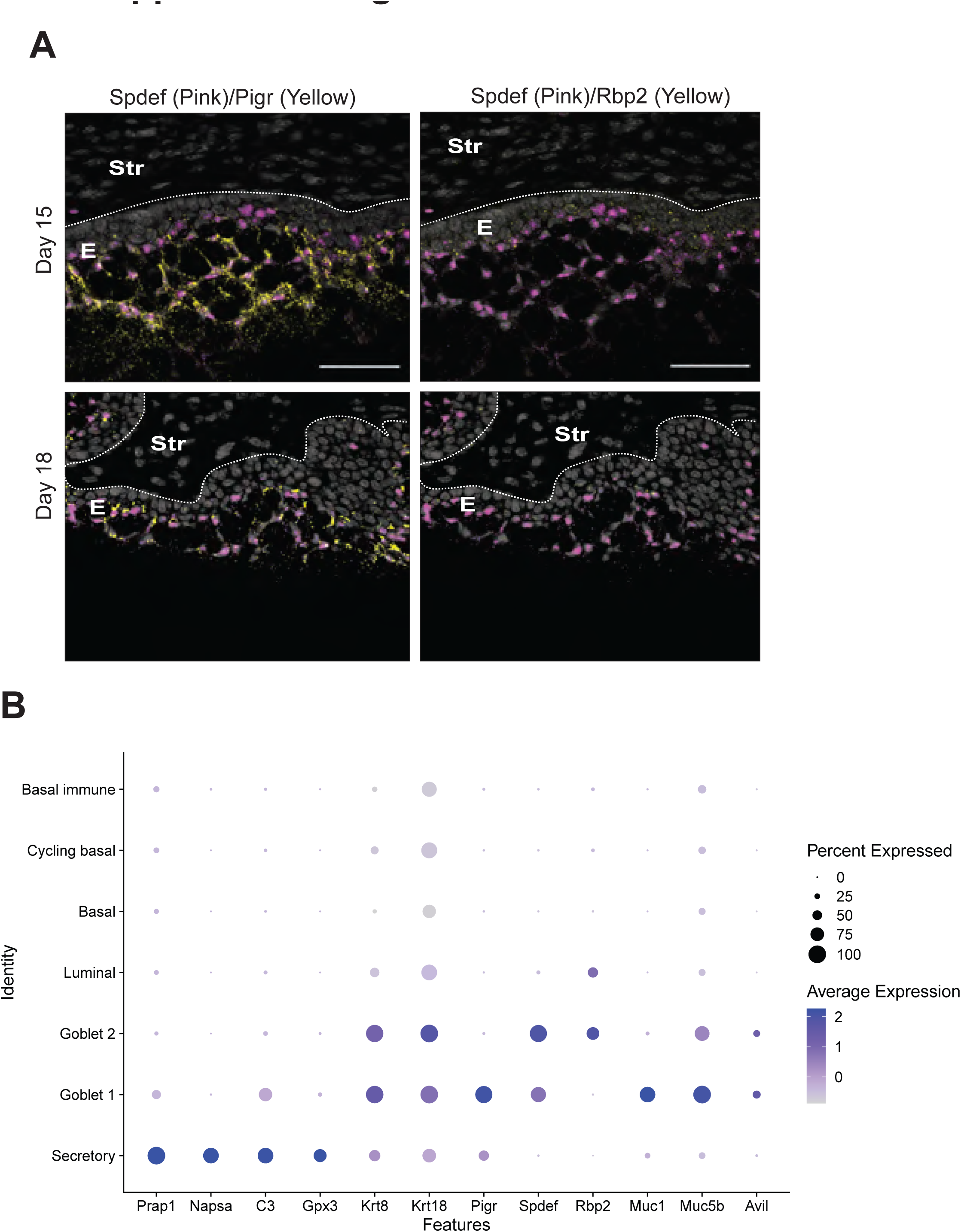
**A.** Spatial transcript analysis for Spdef (pink) with Pigr (yellow) (left panel) or Rbp2 (yellow) (right panel) in mouse endocervical epithelia from GD15 and GD18. DAPI (gray)-nuclei. E: Epithelia; Str: Stroma. Scale bar 50 µm, objective lens 40x. Representative images from two independent samples per group. B. Dot plot shows the expression of genes unique to the minor secretory cluster found in pregnancy time points. Although it shares a similar keratin expression to the secretory clusters, it does not express secretory gene markers.

**Figure S9.**
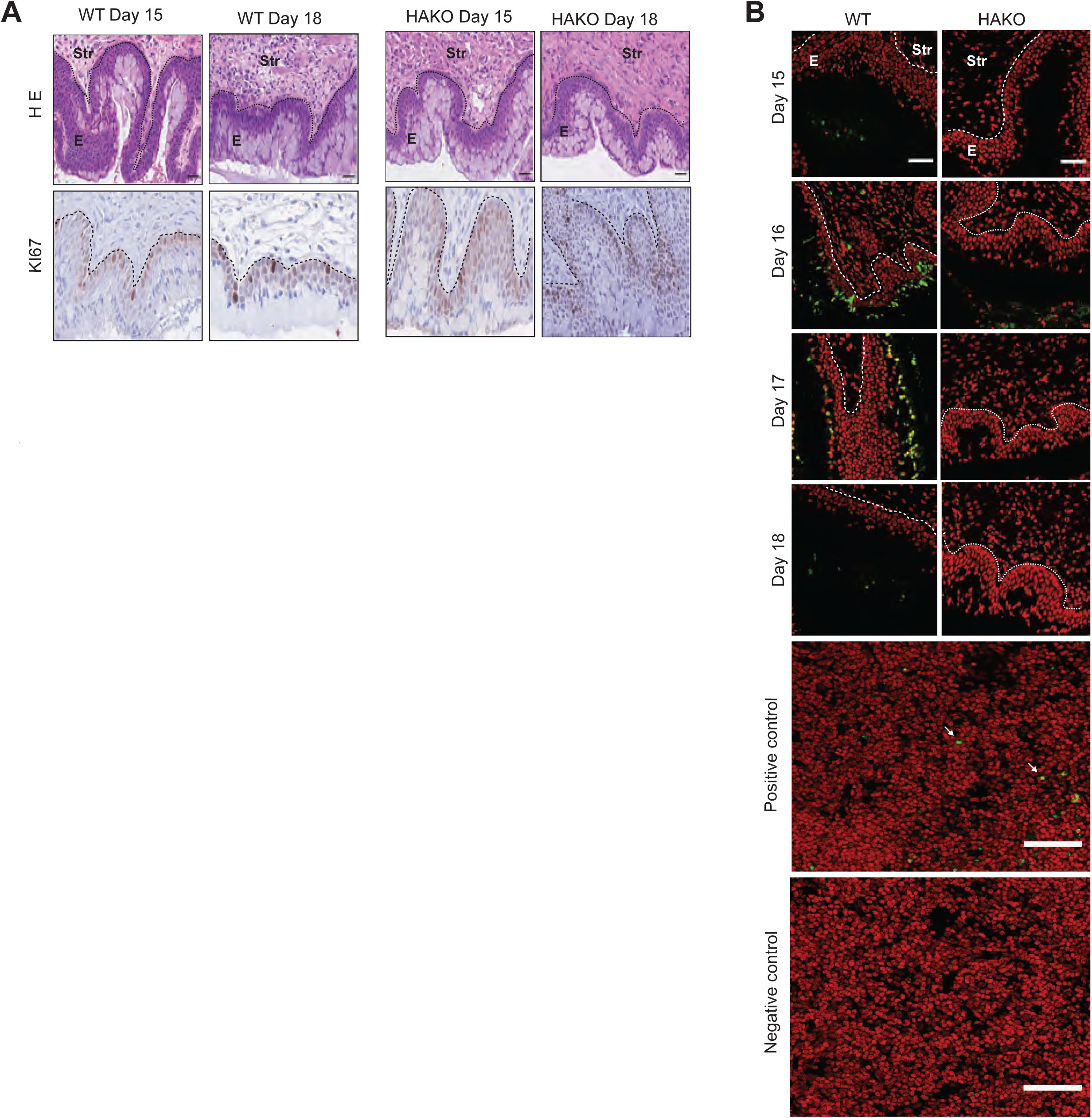
**A.** Comparison of H&E histology and Immunostaining for Ki67 (brown) identify proliferating cells of endocervical epithelia in WT and HAKO at GD15 and GD18. The representative sections are shown at 60x magnification from three independent samples per group. E: Epithelia; Str: Stroma. Scale bar 50 µm B. Comparison of TUNEL staining in endocervical epithelium WT and HAKO at GD15, GD16, GD17 and GD18. Thymus from juvenile wild-type mice was used as positive (with terminal transferase) and negative (without transferase) control. The green cells indicate the TUNEL-positive apoptotic cells (white arrow) while red is nuclei. The representative sections are shown at 20x magnification. Epi: Epithelia; Str: Stroma. Scale bar 50 µm

**Figure S10.**
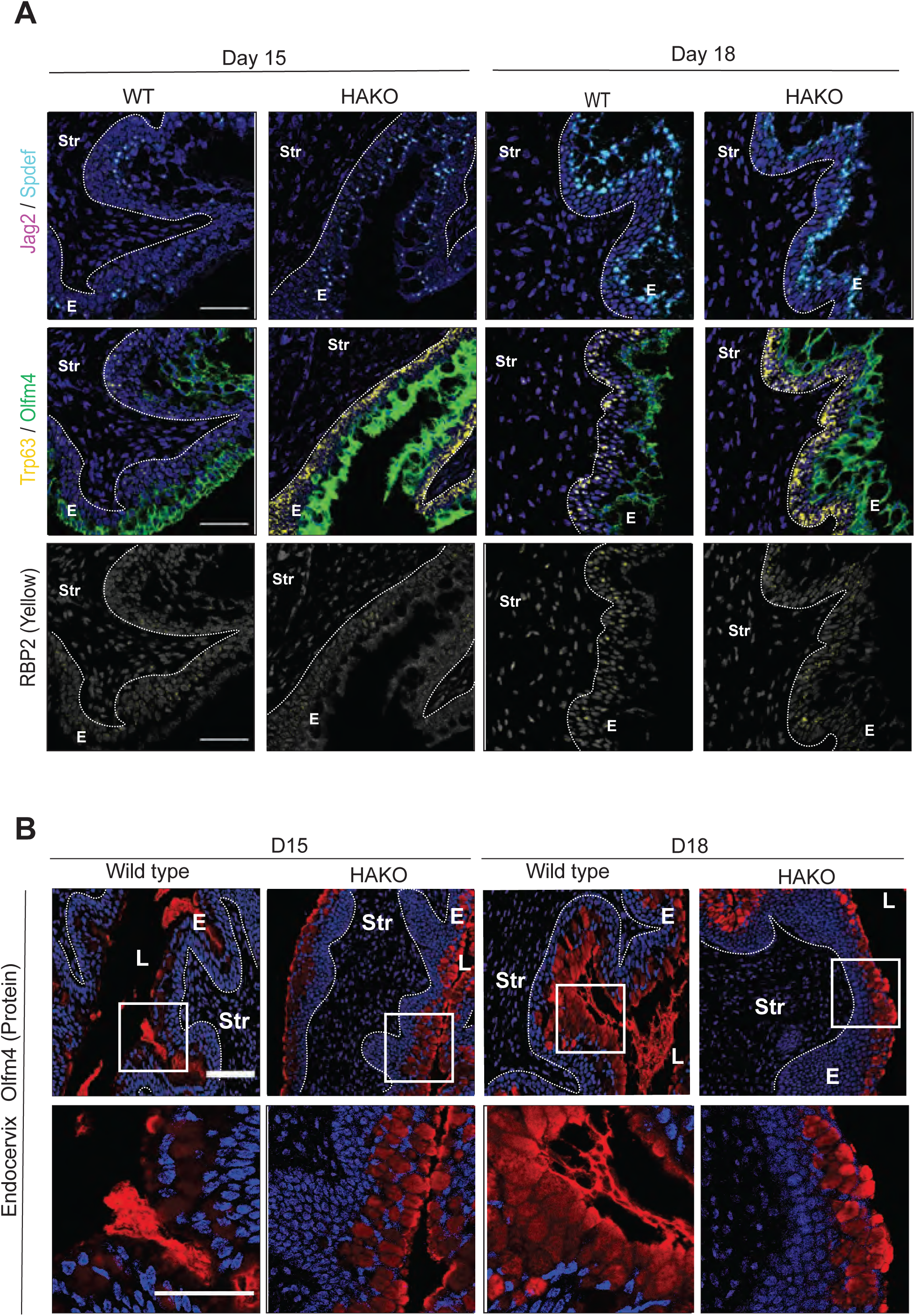
**A.** Comparison of spatial and temporal pattern of epithelial cell markers in WT and HAKO at GD15 and GD18. Fluorescent RNAscope staining for basal and luminal (Jag2, Trp63), and goblet (Spdef, Olfm4, Rbp2) transcripts. Top panel: Jag2 (pink) and Spdef (blue). Middle panel: Trp63 (yellow) and Olfm4 (green). Last panel: Rbp2 (yellow). DAPI stains nuclei (blue or gray). E: Epithelia; Str: Stroma. Scale bar 50 µm, objective lens 40x. Representative images from three independent samples per each group. B. Endocervical epithelia expressing Olfm4 protein (red) in the HAKO on GD15 and GD18 compared to wildtype by IF staining at 20x magnification (Top panel). Images boxed in white are shown at 60x magnification in the bottom panel. DAPI stains nuclei (blue). E: Epithelia; Str: Stroma. Scale bar 50 µm (top and bottom panel). Representative images from three independent samples per group.

**Figure S11.**
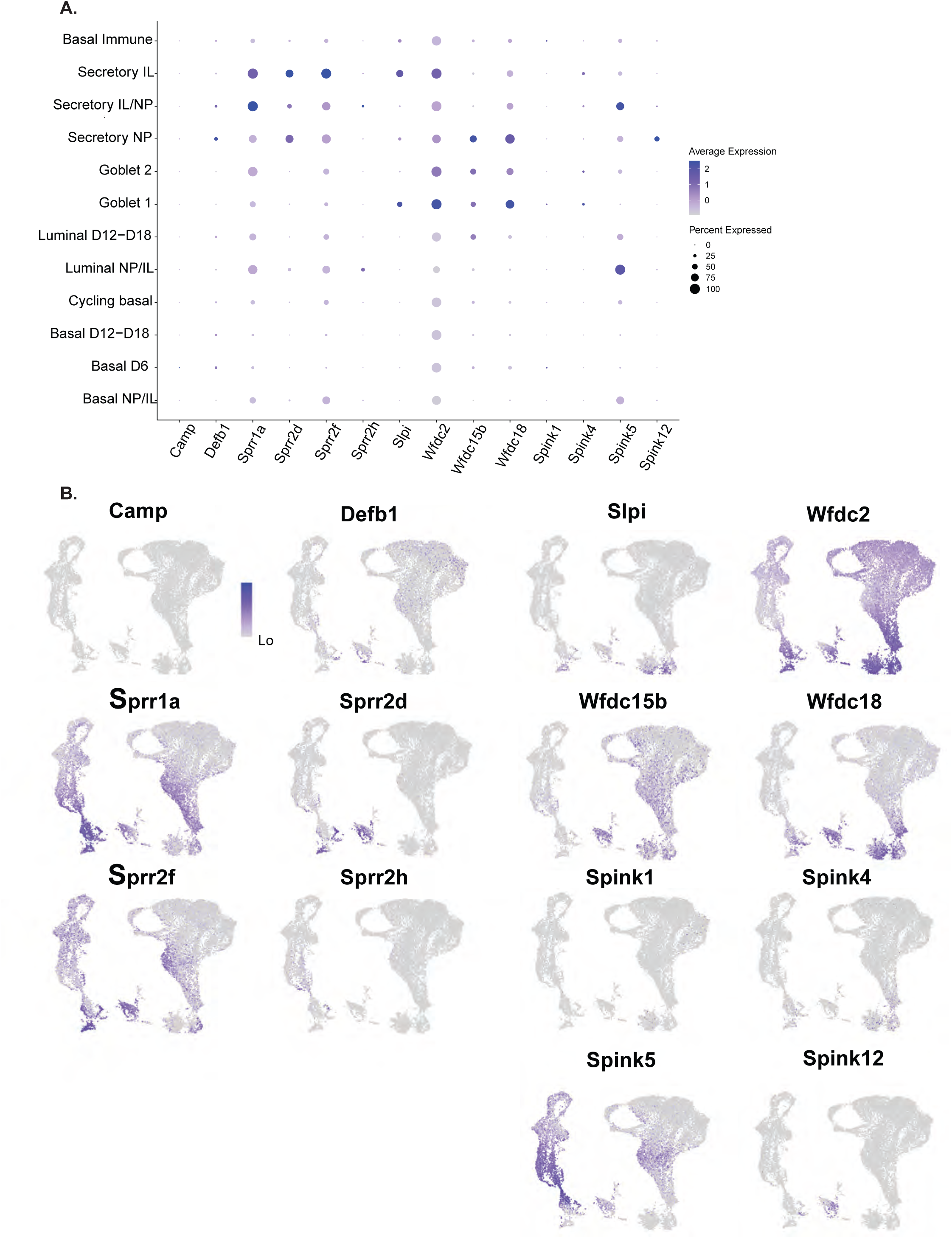
**A.** Dot plot indicating the expression of antimicrobial genes and protease inhibitors in the cervical epithelia. B. Feature plots indicating the expression of antimicrobial genes and protease inhibitors in the cervical epithelia.

## SUPPLEMENTAL TABLE LEGENDS

**Table S1: Sequencing statistics** Includes the following data on scRNA-seq results from each time point collected: number of reads, number of cells captured, and number of cells used for analysis after filtering. Filtering steps removed the following types of cells: doublets, cells with low gene counts, and cells with a high percent of mitochondrial reads.

**Table S2: Differentially expressed genes** Includes the list of differentially expressed genes from 1) the clustering of all cell types from all time points, 2) the clustering of epithelial cells only from all time points, 3) the clustering of epithelial cells only from the HA KO and WT D15/D18 analysis. FindAllMarkers was the Seurat command used to generate these lists.

